# Zfp697 is an RNA-binding protein that regulates skeletal muscle inflammation and regeneration

**DOI:** 10.1101/2023.06.12.544338

**Authors:** Jorge C. Correia, Paulo R. Jannig, Maya L. Gosztyla, Igor Cervenka, Serge Ducommun, Stine M. Præstholm, Kyle Dumont, Zhengye Liu, Qishan Liang, Daniel Edsgärd, Olof Emanuelsson, Paul Gregorevic, Håkan Westerblad, Tomas Venckunas, Marius Brazaitis, Sigitas Kamandulis, Johanna T. Lanner, Gene W. Yeo, Jorge L. Ruas

**Author notes:** **Corresponding author:** Jorge L. Ruas, Molecular and Cellular Exercise Physiology, Department of Physiology and Pharmacology, Biomedicum, Karolinska Institutet, 171 77 Stockholm, Sweden.

## Abstract

Muscular atrophy is a mortality risk factor that happens with disuse, chronic disease, and aging. Recovery from atrophy requires changes in several cell types including muscle fibers, and satellite and immune cells. Here we show that Zfp697/ZNF697 is a damage-induced regulator of muscle regeneration, during which its expression is transiently elevated. Conversely, sustained Zfp697 expression in mouse muscle leads to a gene expression signature of chemokine secretion, immune cell recruitment, and extracellular matrix remodeling. Myofiber-specific Zfp697 ablation hinders the inflammatory and regenerative response to muscle injury, compromising functional recovery. We uncover Zfp697 as an essential interferon gamma mediator in muscle cells, interacting primarily with ncRNAs such as the pro-regenerative miR-206. In sum, we identify Zfp697 as an integrator of cell-cell communication necessary for tissue regeneration.

**One Sentence Summary:** Zfp697 is necessary for interferon gamma signaling and muscle regeneration.

## Introduction

Skeletal muscle is fundamental for breathing, posture, and locomotion and plays a crucial role in the regulation of energy metabolism (*1*). Skeletal muscle mass and function are associated with overall quality of life and health and are inversely correlated with the risk of death. Depending on use and disuse muscle can change its mass, metabolism, and fiber size and type (*2*). Both muscle regeneration and hypertrophy depend on changes in several cell types that include muscle fibers, satellite cells, fibro-adipogenic progenitors and immune cells (*3, 4*). Perturbations in any of these components will affect functional recovery. Indeed, it has been shown that failure to recruit immune cells such as neutrophils, macrophages, and T cells compromises muscle repair and growth (*5, 6*). Accordingly, several cytokines and chemokines have been involved in mediating the cell-cell communication processes necessary for tissue repair. Lastly, remodeling the extracellular matrix (ECM) is necessary to create the right tissue environment and structure to support and transduce force generated by regenerating fibers or as they become larger and stronger. As dependent as muscle regeneration and hypertrophy are on these inflammatory and remodeling processes, failure to resolve them results in tissue fibrosis and progressive loss of function (*7*). This is often seen in muscular dystrophies, where constant tissue injury created by fiber fragility promotes exacerbated immune cell infiltration, permanent tissue inflammation, and excessive ECM deposition and remodeling (*8*). Understanding how these processes are regulated has important implications for clinical and space medicine and will pave the way for the development of therapies for muscle injury, trauma, recovery from intense exercise, and muscle genetic diseases. Here, we uncover the role of the previously uncharacterized gene and protein zinc finger protein 697 (Zfp697 in mice and ZNF697 in humans) as a central regulator of muscle inflammation, regeneration, and compensatory hypertrophy. We show that Zfp697 expression is transiently increased in situations of intense exercise, muscle injury or recovery from atrophy and promotes the expression of genes involved in immune cell recruitment, satellite cell activation, and ECM remodeling. Loss of Zfp697 expression in the muscle fiber severely compromises muscle repair, force recovery, and function, whereas prolonged Zfp697 expression leads to tissue inflammation and fibrosis. In addition, we explore the mechanism of Zfp697 action and regulation by upstream stimuli and identify a critical role in transducing the regenerative actions of interferon gamma (IFN*γ*) in muscle.

### Zfp697 expression is induced during skeletal muscle recovery from atrophy or injury

To identify molecular mechanisms that control skeletal muscle transition from atrophy to compensatory regeneration and hypertrophy, we used a mouse model of hindlimb unloading/reloading (Fig. 1A) (*9, 10*). This protocol results in approximately 30% reduction in gastrocnemius mass after 10 days of unloading, which requires more than 7 days of reloading to recover to control levels (Fig. 1B). To identify gene regulatory events taking place upon muscle reloading, we performed global gene expression analysis by RNA-sequencing (RNA-seq) of gastrocnemius muscle collected from control mice, mice at 10 days of unloading-induced muscle atrophy (hindlimb unloading) and from mice returned to normal cage activity for 1 day (reloading) (Fig. 1C and fig. S1A; Data S1).

**Fig. 1.**
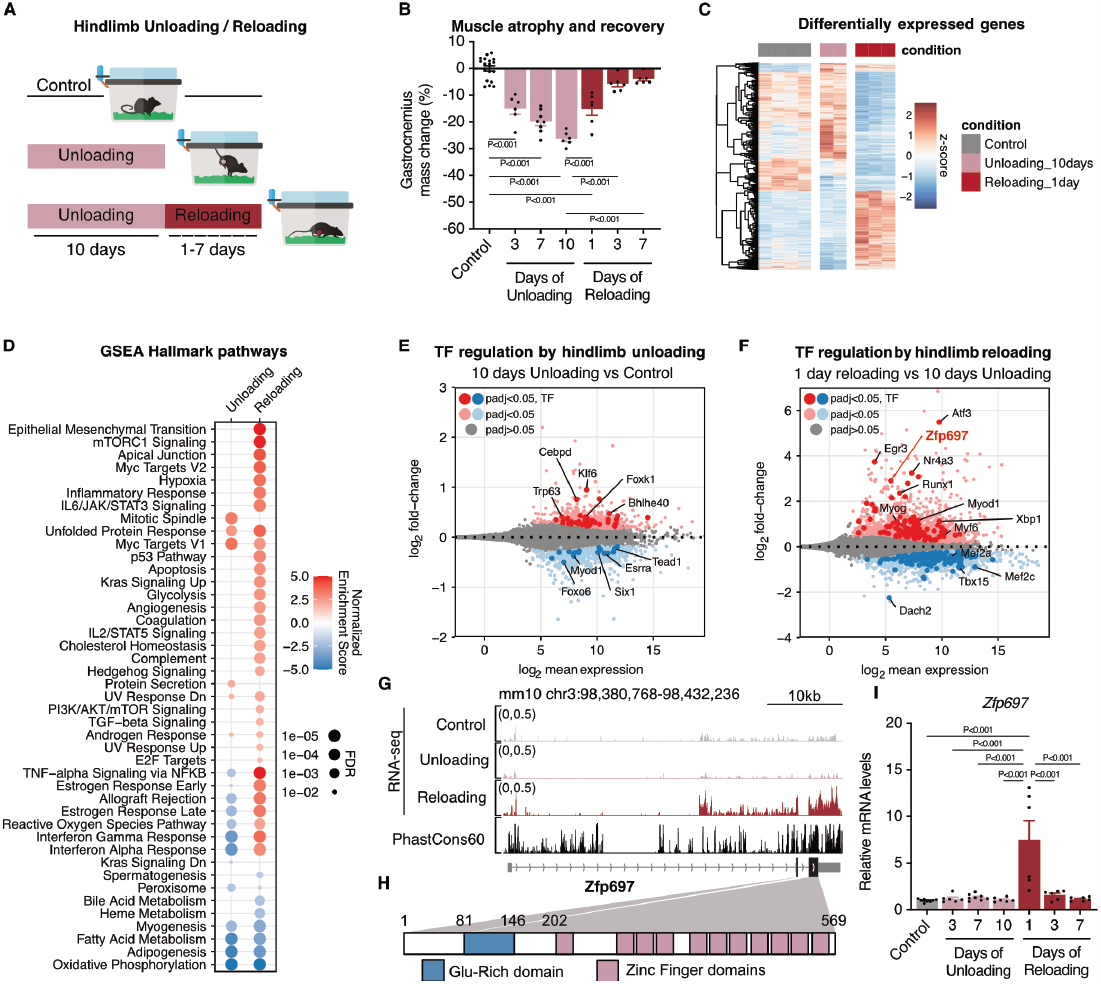
Zfp697 expression increases during muscle recovery from atrophy. (**A**) Overview of the mouse hindlimb unloading and reloading protocol. (**B**) Change in mouse gastrocnemius mass (normalized by tibia length) during hindlimb unloading and reloading (n = 6-18). One-way ANOVA with Tukey’s multiple comparisons test. (**C**) RNA-seq of mouse gastrocnemius muscle after 10 days of hindlimb unloading, 10 days of unloading followed by 1 day of reloading, and controls (n = 2-4 per condition). Heatmap highlights genes differentially expressed between unloading compared with control mice, and reloading compared with unloading. (**D**) Gene set enrichment analysis (GSEA) for hallmark pathways in mouse gastrocnemius after hindlimb unloading and reloading. FDR, false discovery rate. (**E-F**) MA plots of gene expression levels in mouse gastrocnemius after 10 days of unloading compared with control, and 1 day of reloading compared with 10 days of unloading. Red dots indicate genes with significantly increased expression, while blue dots indicate genes with significantly decreased expression (padj < 0.05). Known and putative transcriptional factors are highlighted. TF, transcription factor; padj, adjusted p-value. (**G**) Genome browser tracks of the mouse *Zfp697* gene highlighting RNA-seq data for control, unloaded and reloaded gastrocnemius, and PhastCons60 vertebrate conservation score. (**H**) Schematic representation of the Zfp697 protein and its predicted glutamine-rich and zinc finger structural domains. The numbers shown refer to amino acid positions in the mouse Zfp697 protein. (**I**) *Zfp697* gene expression in mouse gastrocnemius during hindlimb unloading and reloading (n = 6-8). One-way ANOVA with Tukey’s multiple comparisons test. Data represent mean values and error bars represent SEM.

Gene set enrichment analysis (GSEA) of “unloading vs control” and “reloading vs unloading”, performed with hallmark gene sets (*11*), revealed in common between both groups a repression of oxidative metabolism and myogenesis-related pathways and an induction on unfolded protein response (Fig. 1D; Data S2). Among the pathways repressed with unloading but induced with reloading we found NF-kB and IFN alpha and gamma signaling. Unique to the early reloading set were pathways related to epithelial-mesenchymal transition, mTORC signaling, hypoxia and angiogenesis, glycolysis, and TGF-β signaling (Fig. 1D). To identify potential upstream regulators of genes involved in those different biological processes, we overlapped our RNA-seq data with a list of known and predicted transcription factors (TFs) (*12*). With this approach we identified several well-known regulators of muscle stress response (Atf3, Xbp1) (*13*), metabolism (Nr4a3) (*14*), and regeneration (Runx1, Mef2c) (*15*) (Fig. 1, E and F). Among the transcripts increased during the reloading phase we found the previously uncharacterized Zfp697 (which is annotated in Lambert et al., 2018 as a putative TF) (Fig. 1F). Since Zfps are a group of versatile proteins with diverse biological activities and mechanisms of action, that includes several TFs (*16*), we decided to investigate the role of Zfp697 in skeletal muscle. Zfp697 is a highly conserved gene encoding a 569 amino acid protein with an N-terminal glutamine-rich domain, followed by eleven C2H2 zinc finger domains (Fig. 1, G and H). qRT-PCR analysis of muscle samples from the unloading/reloading time course shown in Fig. 1, A and B, confirmed that although Zfp697 levels don’t change during the unloading period, they increase 7.5-fold after 1 day of reloading, and quickly return to baseline at day 3 (Fig. 1I). In a similar fashion, ZNF697 was also increased in human skeletal muscle upon ambulatory reloading following limb unloading (fig. S1, B and C) (*17*). Interestingly, analysis of muscle RNA-seq data from a unloading/reloading protocol using adult vs old mice (*18*), revealed a similar reloading-induced increase in Zfp697 levels in adult mice, that was lost in the aged group (where it is already elevated in controls) (Fig. S1D). Additional analysis of skeletal muscle RNA from different mouse models or interventions revealed that Zfp697 is also transiently elevated in response to β_2_-adrenergic receptor (β_2_-AR) agonism (fig. S2A), acute sub-maximal exercise (fig. S2B), muscle injury (fig. S2C), and lipopolysaccharide administration (LPS, fig. S2D). In human skeletal muscle, ZNF697 expression was also transiently increased following aerobic, resistance, or high intensity interval training (fig. S2, E and F). Interestingly, Zfp697 expression remained elevated in situations of unresolved muscle damage such as muscular dystrophy (fig. S2G) (*19*) or cancer-related cachexia (fig. S2H) (*20*). In agreement with the data shown in fig. S1D, we determined that Zfp697 expression in mouse muscle progressively increases with age (fig. S2I) and is strongly increased in a model of denervation (fig. S2J). ZNF697 expression was also higher in muscle of type 2 diabetic patients, following exercise (fig. S2K) (*21*).

### Zfp697 mediates the interferon gamma response in muscle cells

To determine which cell types within the muscle tissue express Zfp697 we queried the publicly available scRNA-seq resources MyoAtlas (*22*) and datasets for regenerating skeletal muscle stromal vascular fraction (i.e. mononuclear cells) (*23*). These analyses revealed that at baseline Zfp697 is expressed in mature fibers in both tibialis anterior (TA) soleus muscles (fig. S3A and S3B). Both muscles show Zfp697 expression in other cell types most notably in soleus fibro-adipogenic precursors (FAPs) and endothelial cells (fig. S3B). Non-fiber expression can also be seen in mononuclear cells in regenerating muscle (fig. S3C).

To begin uncovering the biological role of Zfp697 we used recombinant adenovirus to increase or silence Zfp697 expression in mouse primary myotubes (vs a GFP or scrambled shRNA control, respectively) (Fig. 2A). Initial analysis by qRT-PCR focused on some of the genes highlighted in the gastrocnemius reloading RNA-seq analysis (Fig. 1D), namely cytokines and chemokines. With this approach we determined that Zfp697 expression in myotubes positively correlated with the mRNA expression for the analyzed cytokines and chemokines, in both gain-and loss-of-function experiments (Fig. 2B). To better understand the overall effects of Zfp697 in myotubes, we performed RNA-seq under the gain-of-function experimental conditions (fig. S4A; Data S3). GSEA for hallmark pathways highlighted, amongst others, IFN response, JAK/STAT signaling, and inflammatory response (Fig. 2C; Data S4) which largely overlapped with pathways observed for the reloaded gastrocnemius muscle (Fig. 1D). In particular, the IFN*γ* response pathway (Fig. 2C) and its main target genes (Fig. 2D) were consistently highlighted. Moreover, GSEA for cellular component pathways suggested the induction of transcriptional programs related to ECM remodeling, vesicle trafficking and secretory apparatus (fig. S4B). Analysis of distant regulatory elements (DiRE) (*24*) predicted to mediate the observed gene expression patterns, revealed that genes induced by Zfp697 are also potentially regulated by the transcription factor families STAT and SMAD (fig. S4C). Interestingly, there are several evolutionarily conserved STAT binding sites within the proximal promoter for the *Zfp697* gene (fig. S4D), which proved to be inducible in myotubes by IFN*γ* treatment but not by TNFα (fig. S4E). Together, these data and the fact that IFN*γ* has been implicated in muscle regeneration (*25, 26*), prompted us to investigate whether Zfp697 is part of the IFN*γ* signal transduction pathway. We next treated mouse primary myotubes with recombinant IFN*γ* after silencing Zfp697 expression (vs a scrambled control shRNA). Although *Zfp697* transcription was only modestly induced by IFN*γ* treatment, it was required for the basal expression of several chemokines and known IFN*γ* target genes and proved to be strikingly necessary for IFN*γ* response (Fig. 2. E and F). When comparing genes regulated by Zfp697 expression in primary myotubes with genes differentially expressed in the reloaded mouse gastrocnemius, shared genes with increased expression (535 genes) clustered around pathways of cell adhesion, immune cell recruitment, and cytokine-mediated cell-cell communication, whereas genes with commonly decreased expression highlighted different metabolic pathways (fig. S4G; Data S5). Focusing this analysis specifically on IFN*γ* target genes, we observed a significant overlap between those increased by ectopic Zfp697 expression in myotubes and those increased during the early reloading stage of the unloading/reloading protocol (when Zfp697 itself is increased) (fig. S4H).

**Fig. 2.**
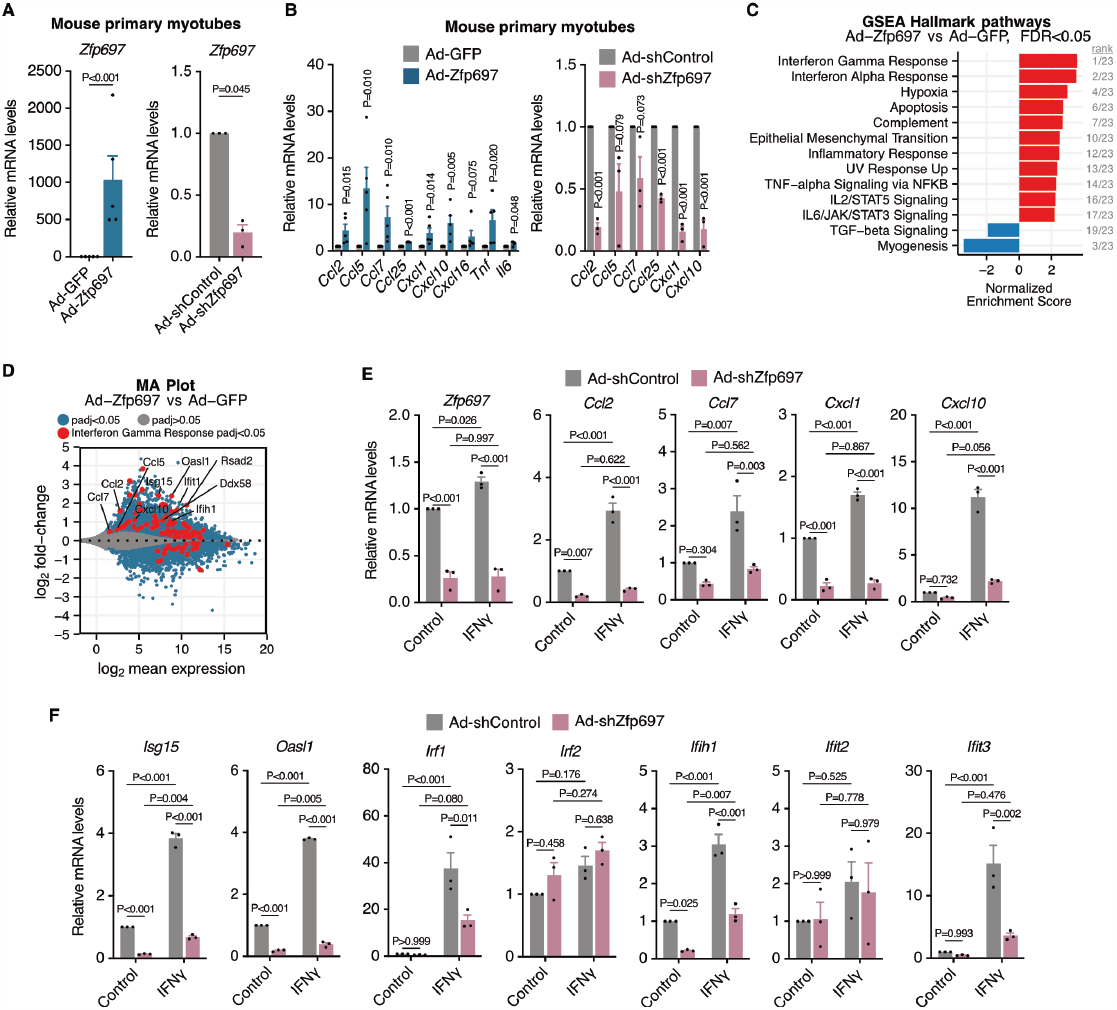
Zfp697 is necessary for IFN*γ* signal transduction in myotubes. (**A**) *Zfp697* gene expression in mouse primary myotubes with Zfp697 gain- or loss-of-function. For gain-of-function experiments, fully differentiated myotubes were transduced with adenovirus expressing GFP alone (Ad-GFP) or together with Zfp697 (Ad-Zfp697; n = 4 independent experiments). For loss-of-function, cells were transduced with adenovirus expressing Zfp697-specific (Ad-shZfp697) or scrambled control shRNAs (Ad-shControl; n = 3 independent experiments). Two-tailed Student’s paired t test versus respective control. (**B**) Gene expression analysis of chemokines in mouse primary myotubes with Zfp697 gain- or loss-of-function. Two-tailed Student’s paired t test versus respective control. (**C**) Gene set enrichment analysis for hallmark pathways in mouse primary myotubes overexpressing Zfp697. FDR, false discovery rate. (**D**) MA plot showing mean gene expression versus log_2_ fold-change in mouse primary myotubes overexpressing Zfp697. Differentially expressed genes are shown in blue (padj < 0.05). Interferon gamma response genes are highlighted in red. padj, adjusted p-value. (**E-F**) Gene expression analysis of *Zfp697*, chemokines and interferon gamma (IFN*γ*) response genes in mouse primary myotubes after Zfp697 loss-of-function and treated with IFN*γ* or control (20 ng/mL for 8 hours; n = 3 independent experiments). Two-way ANOVA with Tukey’s multiple comparisons test. Data represent mean values and error bars represent SEM.

### Zfp697 is an RNA-binding protein that targets lncRNAs and miRNAs

Since Zfp697 is mostly composed of zinc finger domains, we first thought it could function as a DNA-binding TF. To test that possibility, we performed chromatin immunoprecipitation followed by sequencing (ChIP-seq) in mouse primary myotubes expressing Flag-Zfp697 or a GFP control. Surprisingly, the results from these experiments revealed very few chromatin regions associated with Flag-Zfp697 (fig. S5A). Although these data do not conclusively prove that Zfp697 does not bind DNA, they suggest that Zfp697 might not work primarily as a TF. In support of this notion, we observed that purified GST-fused Zfp697 can interact directly with RNA (Fig. 3A). This interaction relied on the zinc finger domains, since a truncated mutant lacking the zinc finger domains bound RNA to the same level as GST alone (Fig. 3A).

**Fig. 3.**
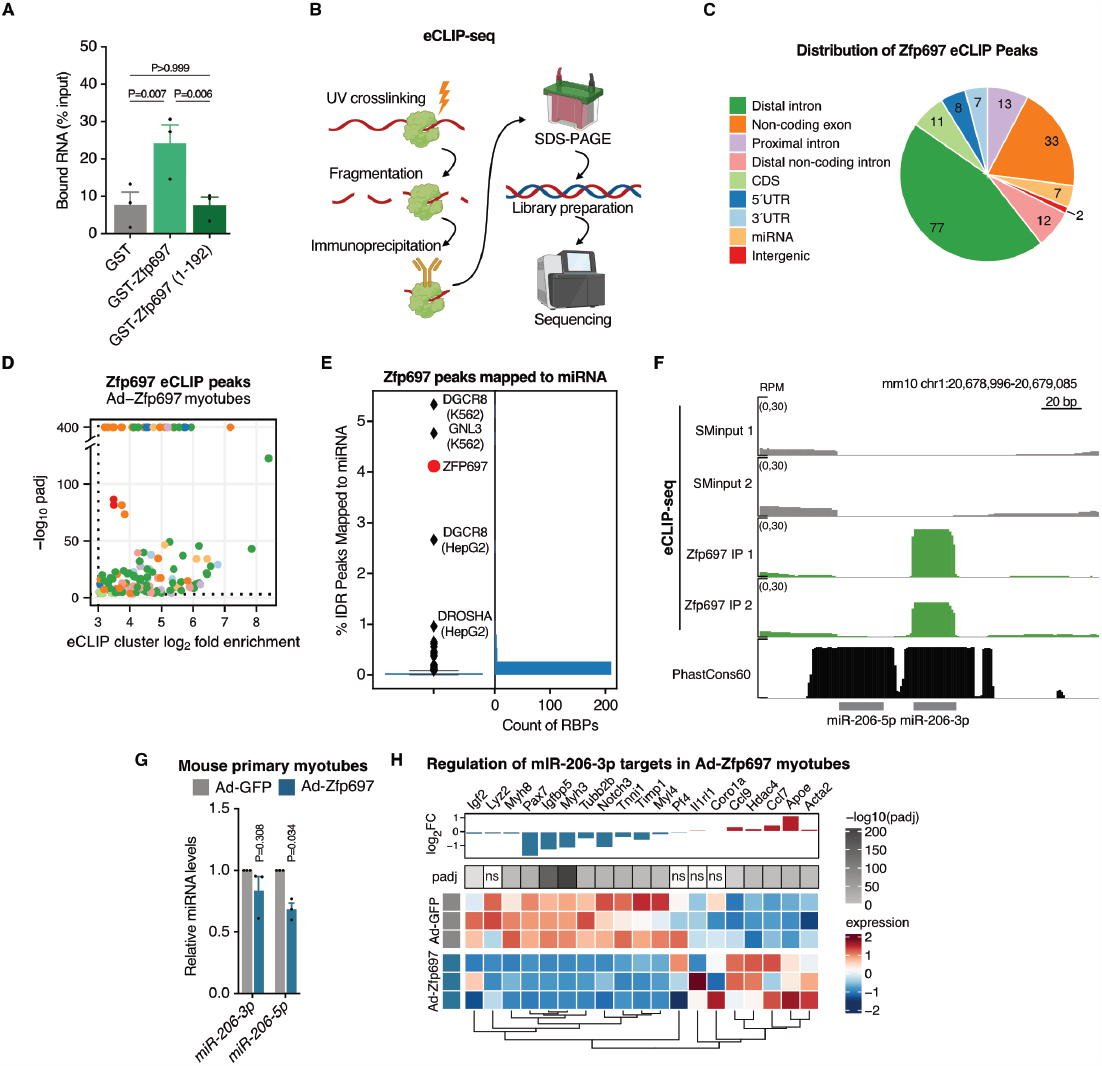
Zfp697 is an RNA-binding protein that interacts with ncRNAs and miRNAs. (**A**) In vitro RNA-binding assay performed with GST alone (control), full-length Zfp697, and Zfp697 N-terminus (aa 1-192) lacking all the zinc finger motifs. Results are displayed as percentage of input recovered (n = 3-4). One-way ANOVA with Fisher’s LSD. (**B**) Overview of the enhanced UV crosslinking followed by immunoprecipitation (eCLIP) sequencing protocol. **(C-D)** Volcano plot (bottom left panel) and distribution (top right panel) of significantly enriched Zfp697 eCLIP peaks in mouse primary myotubes overexpressing Zfp697. Cutoffs of eCLIP IP/SMInput log_2_ fold enrichment and -log_10_ padj are denoted by dashed lines (padj < 0.001, fold enrichment > 8, IDR < 0.01). Color coding represents location of Zfp697 binding sites across genic regions. padj, adjusted p-value; IDR, Irreproducible Discovery Rate. (**E**) Proportion of significantly enriched binding sites mapped to miRNA for Zfp697 (red) compared to other RNA binding proteins (RBPs) with eCLIP data in ENCODE 3. (**F**) Genome browser tracks showing Zfp697 binding site at miR-206-3p. PhastCons60 represents vertebrate conservation score. SMInput, size-matched input; RPM, reads per million. (**G**) Expression of miR-206-3p and miR-206-5p in mouse primary myotubes overexpressing Zfp697 (n = 3 independent experiments). Two-tailed Student’s paired t test versus Ad-GFP. (**H**) Expression heatmap and log_2_ fold-change of miR-206-3p target genes in mouse primary myotubes overexpressing Zfp697 determined by RNA-seq. padj, adjusted p-value; ns, not significant.

To further investigate the RNA-binding properties of Zfp697 and identify Zfp697-bound RNAs, we performed enhanced crosslinking and immunoprecipitation (eCLIP) using the same primary myotube experimental setup as for the ChIP-seq experiments but analyzed by eCLIP followed by high-throughput sequencing (eCLIP-seq, Fig. 3B and fig. S5B) (*27*). With this approach we identified 170 Zfp697-associated eCLIP peaks (Fig. 3, C and D; Data S6), corresponding to 98 unique genes, that were significantly enriched compared to a size-matched control (p < 1e-3, FC > 8). Nearly half of the significant peaks mapped to RNAs that originate from distal intronic regions (Fig. 3C). The median length of these distal intronic peaks was 26 bp, which is close to the typical size of miRNAs (22 bp), suggesting that some of these peaks could represent unannotated miRNAs (fig. S5C). Additionally, approximately 4% of the peaks mapped to annotated miRNAs. To normalize these results for the different average lengths of each type of transcript feature, we applied a region-based CLIP enrichment workflow (https://github.com/YeoLab/region_based_CLIP_enrichment). The length-normalized analysis showed that the proportion of peaks mapping to miRNAs is enriched 2.3-fold compared to a null distribution. Indeed, when compared to eCLIP results from the ENCODE 3 dataset (n=223 eCLIP experiments) (*27*), the proportion of Zfp697 peaks mapping to miRNAs is higher than 99% of other datasets, even exceeding the known miRNA processing factor DROSHA (Fig. 3E). Compared to ENCODE 3, miRNA was the most highly over-represented transcript type (fig. S5D). Other genomic locations showing high association with Zfp697 were non-coding exons, with the least represented locations being coding sequences (fig. S5D). Interestingly, one of the top hits among Zfp697 miRNA targets was miR-206-3p, which has been previously shown to promote skeletal muscle regeneration (Fig. 3F) (*28, 29*). In line with that observation, we determined that myotubes overexpressing Zfp697 exhibit reduced abundance of the miR-206-5p passenger strand, suggesting Zfp697 promotes the usage of miR-206-3p as the mature strand (Fig. 3G) (*30*). In agreement, analysis of the mRNA abundance for miR-206-3p targets previously reported to be downstream of its regenerative effects in muscle (*28*), showed the expected reduction in Zfp697-expressing myotubes, although not for all genes (Fig. 3H).

### Myofiber-specific Zfp697 knockout compromises muscle recovery from atrophy and injury

To test the effects of Zfp697 expression in vivo, we used adenovirus-mediated gene delivery by intramuscular injection in SCID mice (vs a GFP control in the contralateral limb; fig. S6A). This resulted in a mild but sustained overexpression of Zfp697 until 7 days post-injection (fig. S6B). Targeted qRT-PCR analysis of some of the pathways highlighted by the previous RNA-seq data sets, revealed that muscles transduced with the Zfp697-encoding adenovirus (vs the GFP transduced contralateral limb) had higher expression levels of several chemokines and their receptors (fig. S6, C and D), genes related to IFN signaling (fig. S6E), immune cell markers (fig. S6F), and ECM remodeling genes (fig. S6G). In line with these gene expression patterns, we could detect signs of tissue fibrosis by immunohistochemical analysis of the transduced muscle (fig. S6, H and I). Importantly, we also determined that Zfp697/ZNF697 expression is robustly increased in situations of disease with a strong unresolved inflammation and fibrosis component. These included mouse kidney injury and different mouse models of heart failure and dysfunction (fig. S7).

To evaluate the consequences of loss of Zfp697 function in muscle, we generated a myofiber-specific Zfp697 knockout mouse (Zfp697 mKO). This was achieved by creating a mouse line carrying loxP sites flanking the coding sequence within exon3 of Zfp697 (which encodes 86% of the protein, including all zinc finger domains) (Fig. 4A), which was crossed with the Mef2c enhancer/myogenin promoter-Cre line (*31*) (Fig. 4A and fig. S8A). Zfp697 mKOs showed reduced but not absent Zfp697 expression in muscle tissues (reflecting Zfp697 expression in other cell types), and no changes in any other analyzed tissue (fig. S8B).

**Fig. 4.**
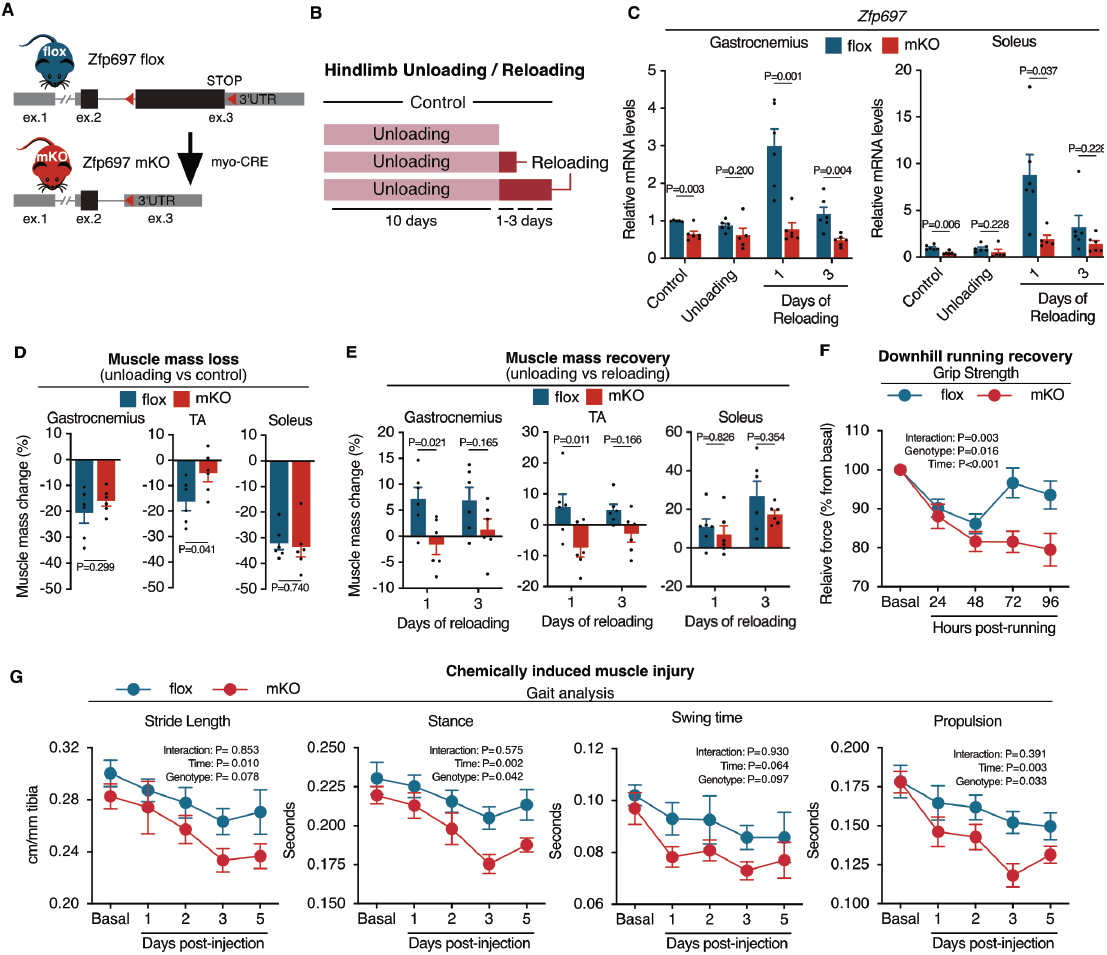
Skeletal muscle-specific Zfp697 knockout blunts recovery from injury. (**A**) Schematic representation of the strategy adopted to generate skeletal-muscle specific *Zfp697* knockout mice (Zfp697 mKO) and flox controls. Red arrows indicate loxP sites location. (**B**) Experimental approach for hindlimb unloading and reloading of Zfp697 mKO and flox littermates. (**C**) *Zfp697* gene expression in the gastrocnemius (left panel) and soleus (right panel) of Zfp697 mKO and flox littermates following hindlimb unloading and reloading. Two-tailed Student’s t test compared with time-matched flox controls. (**D**) Muscle mass change after 10 days of hindlimb unloading comparing Zfp697 mKO and flox littermates with pre-unloading genotype-matched controls (n = 6). Two-tailed Student’s t test. (**E**) Muscle mass change after 1 or 3 days of hindlimb reloading comparing Zfp697 mKO and flox littermates with genotype-matched 10 days of unloaded mice (n = 6). Two-way ANOVA with Šídák’s multiple comparisons test. (**F**) Change in body-weight-normalized grip strength following a single bout of strenuous downhill running in Zfp697-mKO mice and flox littermates (n = 10). Repeated measures two-way ANOVA with Šídák’s multiple comparisons test. (**G**) Gait analysis of Zfp697 mKO mice and floxed littermates (flox, n = 5; mKO, n = 4) at baseline and following chemically induced muscle injury. Cardiotoxin was intramuscularly injected in the gastrocnemius and tibialis anterior muscles of one leg (right) and control solution in the same site of contralateral leg (left). Repeated measures two-way ANOVA. Data represent mean values and error bars represent SEM.

At baseline, no differences were seen between genotypes in terms of body weight, muscle tissue mass, exercise performance, or grip strength (fig. S8, C to F). Zfp697 mKO mice and floxed littermate controls were then subjected to the unloading/reloading protocol. Each genotype was divided into control (no unloading or reloading), 10 days unloading, and two additional groups that were unloaded for 10 days followed by 1 or 3 days of reloading (Fig. 4B). In line with our previous results, *Zfp697* expression was elevated in muscles from floxed controls during reloading (Fig. 4C). This regulation was completely blunted in Zfp697 mKO mice (Fig. 4C), further indicating that, in this context, that response comes mostly from the muscle fibers. Analysis of total body mass and composition during the experimental protocol revealed that Zfp697 mKO mice lost total, lean, and fat mass to a similar extend as floxed littermates, during the unloading phase (fig. S8G). In line with this, changes in muscle mass during the unloading phase were generally comparable between Zfp697 mKOs and floxed littermates except for the TA, that showed reduced wasting in Zfp697 mKO mice (Fig. 4D). Conversely, upon reloading Zfp697 mKOs showed reduced recovery of muscle mass in both gastrocnemius and TA muscle, with no significant differences observed in the soleus (Fig. 4E). In line with these observations, Zfp697 mKO mice had delayed recovery of lean mass, as shown by a significant reduction in lean mass one day after reloading, compared to flox littermates (fig. S8G). The deficit in recovery observed in Zfp697 mKO mice was further evidenced by reduced extensor digitorum longus (EDL) and soleus absolute and specific force 3 days after reloading, especially at higher stimulation frequencies, as evident by significant interaction between genotype and stimulation frequency (fig. S8, H and I). Notably, there were no significant differences in fatigability and recovery (fig. S8, H and I). To further evaluate the implications of myofiber Zfp697 ablation in the functional recovery from muscle injury, we used two further injury models. Firstly, we evaluated the kinetics of muscle force-production recovery after a single bout of downhill running, commonly used approach to elicit exercise-induced muscle damage due to the extensive use of eccentric contractions. Under these conditions, floxed control mice displayed the expected decline in grip strength over the first 48h, followed by a swift recovery to normal values (Fig. 4F). In sharp contrast, Zfp697 mKO did not recover any grip strength by the latest time point analyzed (96h) (Fig. 4F). In addition, we performed intramuscular cardiotoxin (CTX) injections, a myotoxic agent, and monitored functional decline and recovery using gait analysis (Fig. 4G). This assessment revealed a more pronounced decline in various gait parameters in Zfp697 mKO mice, including stride length, stance, and propulsion (Fig. 4G). Together, these data from various models of muscle injury and recovery indicate that Zfp697 ablation in myofibers compromises skeletal muscle regeneration and functional recovery.

### Reloaded Zfp697 mKO muscle fails to activate a regenerative gene program

To complement the functional analysis of the unloading/reloading experiments we performed RNA-seq of the gastrocnemius muscles of the different groups of Zfp697 mKO and controls. This analysis was designed to account for the interaction effect between conditions (control, unloading and reloading) and genotypes. When comparing Zfp697 mKOs and controls for changes in gene expression upon unloading, we saw no differences between genotypes (Fig. 5, A and B and fig. S9A; Data S7) but observed a profound effect on the regenerative response to reloading (Fig. 5, A and C; Data S7). Among the differentially expressed genes, most increased in controls upon reloading but failed to be induced in Zfp697 mKO muscle (Fig. 5B), with a smaller subset changing in the opposite direction. These genes were associated with stress response, protein synthesis, IFN*γ*, and inflammatory responses (Fig. 5, C and D; Data S8), indicating failure of Zfp697 mKO muscle to activate a regenerative response to injury. Muscle regeneration is a complex process that relies on the coordinated action of multiple cell types, many of which are recruited and/or proliferate at the regenerating site. To better understand how Zfp697 ablation in myofibers affected other resident and recruited cell types during reloading-induced regeneration, we deconvoluted our RNA-seq data using CIBERSORTx (*32*) and a previously published single-cell RNA-seq data obtained from mouse regenerating muscle (*23*).

**Fig. 5.**
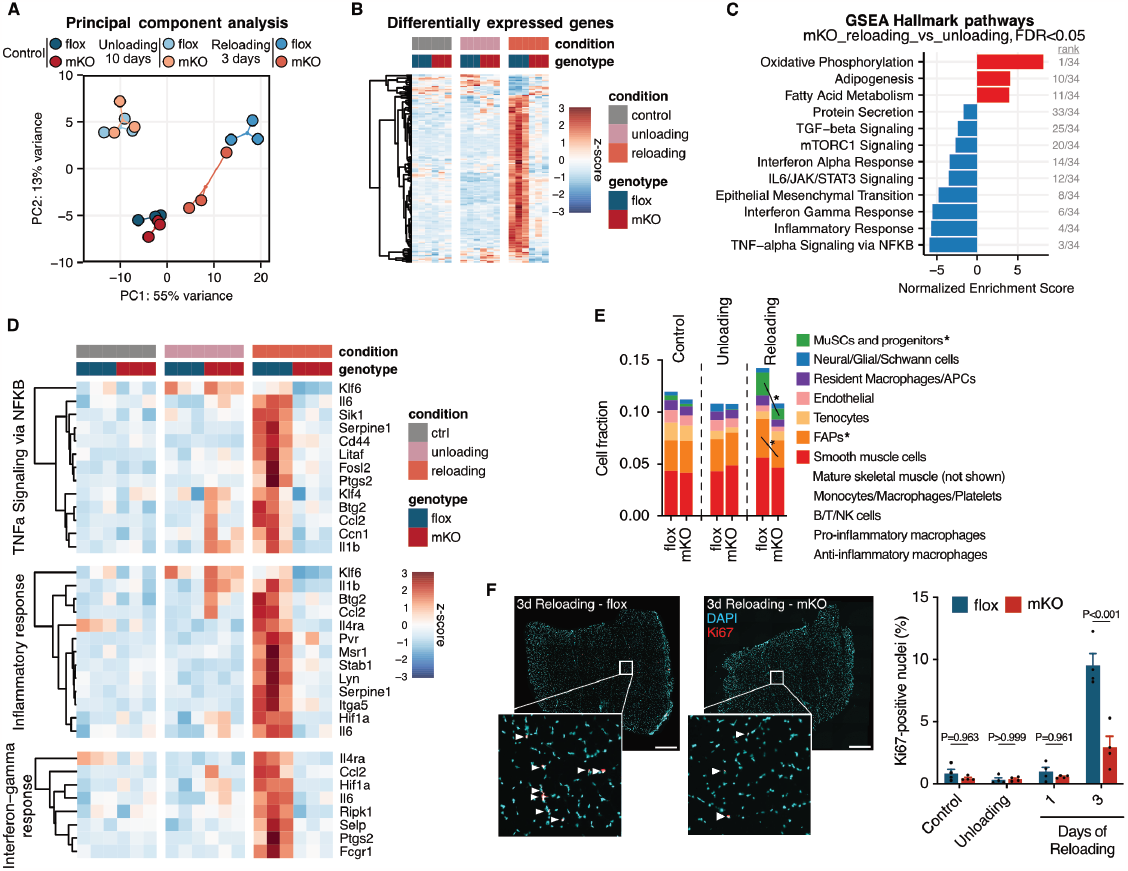
Zfp697 mKOs fail to activate gene expression necessary for regeneration. (**A**) Principal component analysis for RNA-seq performed in the gastrocnemius muscle of control Zfp697 flox and mKO, after 10 days of hindlimb unloading and 3 days of reloading (n = 3 per condition and genotype). (**B**) Heatmap of differentially expressed genes for the interaction effect between conditions and genotypes. (**C**) Gene set enrichment analysis for hallmark pathways accounting for the interaction effect between conditions (control, unloading and reloading) and genotypes (mKO and flox). FDR, false discovery rate. (**D**) Heatmap of genes belonging to inflammatory and interferon gamma response pathways. (**E**) Fraction of muscle-resident cell populations identified by digital cytometry (CIBERSORTx) applied to bulk muscle RNA-seq data from control, hindlimb unloaded and reloaded Zfp697 flox and mKO mice (n = 3 per condition and genotype). Two-way ANOVA with Šídák’s multiple comparisons test. *p<0.05 between indicated cell types. (**F**) Immunostaining for proliferating cells using KI67 in gastrocnemius muscles of Zfp697 flox and mKO and respective quantification (n = 4 per condition and genotype). Two-way ANOVA with Fisher’s LSD. P values represent comparison with time-matched flox controls. Scale bar = 500 μm.

This analysis showed that after 3 days of reloading, the fraction of fibro-adipogenic precursors (FAPs) and muscle stem cells (satellite cells, MuSCs) and progenitors expanded in control muscle, but this response was blunted in Zfp697 mKOs (Fig. 5E and fig. S9B). In line with these data, we observed an increase in the number of proliferative KI67-positive cells in control muscles, which was significantly reduced in Zfp697 mKOs (Fig. 5E). Overall, these data indicate that Zfp697 mKO mice failed to activate a regenerative transcriptional program in myofibers during the early stages of recovery from hindlimb unloading, which affected not only the myofibers themselves but also other resident and infiltrating cells.

Together, our data identify Zfp697 as an RNA binding protein (RBP) that preferentially binds ncRNAs, in particular miRNAs, and controls muscle regeneration. In skeletal muscle, Zfp697 is downstream of IFN*γ* signaling, and is necessary for the activation of many of its target genes. Loss of Zfp697 action results in compromised muscle regeneration, whereas sustained expression leads to tissue inflammation and fibrosis. Collectively, these findings uncover Zfp697 as a promising therapeutic candidate for diseases where unresolved tissue inflammation and ECM remodeling are central disease components.

## Discussion

Progressive loss of skeletal muscle mass and function is a major clinical feature of several neuromuscular diseases, is commonly observed patients with chronic disease such as cancer and is a major contributor to frailty and loss of independence during aging. To better understand the molecular determinants of skeletal muscle atrophy and subsequent recovery, we used a mouse model of hindlimb hypokinesia/hypodynamia (*33*) often used to assess the effects of, for example, long-term bedrest and space flight on muscle mass and function. This protocol, commonly referred to as hindlimb unloading/reloading, offers the advantage of monitoring phenotypic and molecular changes during the progression of muscle atrophy and compensatory hypertrophy/regeneration over time, using the same animals. Among prominent transcriptional regulators of myogenesis, inflammation, and stress response (e.g. Myog, MyoD1, Atf3, RunX1), all of which are important components for the regenerative response to damage, we identified Zfp697 as a previously uncharacterized transcript which is elevated during the early stages of hindlimb reloading. We further verified that mouse Zfp697 expression is transiently induced in multiple settings associated with myofiber damage and remodeling, such as chemically-induced injury and intense physical exercise. By analyzing a vast array of published transcriptomics data, we observed a similar expression profile in human skeletal muscle. In line with its expression profile, Zfp697 expression is enriched in a regenerative myonuclear population recently identified in dystrophic mouse muscle (RegMyon) (*34*) and a yet uncharacterized Ampd3^+^ (AMP deaminase 3) myonuclear population in aged mouse muscle (fig. S3A), hypothesized to constitute dysfunctional denervated fibers (*22*). In support of this notion, Zfp697 expression is highly induced in denervated mouse muscle (fig. S2J). All these data position Zfp697 at the center of the regenerative response in several physiological and pathophysiological contexts.

Muscle regeneration is a complex process that relies on the coordinated action of multiple resident and infiltrating cells. In this context, inflammatory signals have emerged as important regenerative signals, implicated not only in the regulation of immune cell behavior but also satellite cell proliferation, myogenesis, FAP activation, ECM remodeling, and myofiber metabolic adaptions (*3, 35*). Here, we show that activation of Zfp697 in the muscle fiber is upstream of that cascade of events. Zfp697 activates a broad inflammatory gene program in cultured myotubes and intact mouse muscle, including a large array of chemokines implicated in immune cell recruitment and muscle regeneration (*35–39*). The transcriptional profile brought about by ectopic Zfp697 expression was particularly enriched for the IFNα and *γ* pathways. Conversely, IFN α and *γ* response pathways were amongst the top pathways associated with genes dysregulated in Zf697 mKO mice upon hindlimb reloading. IFN*γ* signaling is activated in the early stages of skeletal muscle regeneration (which is further reinforced by our transcriptomics data on reloaded muscle) and endogenous IFN*γ* is necessary for efficient skeletal muscle regeneration (*25, 40*). Moreover, the age-related decline in tissue macrophage responses to IFN*γ* contributes to poor regenerative capacity, by impairing satellite cell proliferation and differentiation (*26*). Strikingly, Zfp697 knockdown in myotubes reduced not only the basal expression of interferon stimulated genes, but also strongly blunted the response to exogenous IFN*γ* indicating that Zfp697 is an important mediator of IFN signaling in skeletal muscle cells. Interestingly, genetic variants in the ZNF697 genomic locus have been associated with the response of multiple sclerosis patients to IFNβ treatment (*41*), indicating that the role of Zfp/ZNF697 in IFN signaling is conserved in humans and in multiple cell types.

In line with the transient nature of Zfp697 activation in regenerating skeletal muscle, myofiber-specific *in vivo* loss-of-Zfp697 function (Zfp697 mKO mice), has negligible effects on animals kept under control conditions, or even when they are subjected to hindlimb unloading. Likewise, the transcriptional profile of Zfp697 mKO at rest or following hindlimb unloading was virtually indistinguishable from that of flox littermate controls. This is somewhat surprising (at least to this extent) and does not recapitulate the strong effect of Zfp697 loss-of-function on the expression of chemokines and interferon-stimulated genes in cultured myotubes. It is possible that the higher expression of these transcripts in other cell types masks potential differences in gene expression in myofibers. It is also possible that the effects of Zfp697 are potentiated in regenerating fibers, a phenotype that may be better reflected by differentiated myotubes than by unchallenged mature fibers. In sharp contrast, the regenerative response to damage is severely compromised in Zfp697 mKO mice. Using three different muscle injury-recovery paradigms (unloading/reloading, downhill running, and cardiotoxin) and complementary methods to monitor recovery over time, we were able to determine that myofiber Zfp697 is essential for skeletal muscle repair and functional recovery. Following injury, Zfp697 mKO mice display deficient recovery of muscle mass, myofiber contractile force, grip strength, and normal gait paremeters. At the molecular level, this was accompanied by deficient activation of a regenerative gene program associated with inflammation, ECM remodeling, angiogenesis, and cell proliferation. Furthermore, Zfp697 mKOs show reduced satellite cell and fibro-adipogenic precursor activation upon hindlimb reloading, two cellular processes that are essential for efficient muscle regeneration (*4, 42, 43*).

Whereas transient activation of a regenerative response conducive to immune cell recruitment and ECM remodeling is critical for efficient repair and functional recovery, persistent activation of these pathways can lead to morphologic and metabolic abnormalities, overt fiber damage, and fibrosis. This is a hallmark of neuromuscular diseases, such as dermatomyositis and muscular dystrophy, and a common feature of cancer cachexia, metabolic diseases, and aging (*44, 45*). We found that the levels of Zfp697/ZNF697 are elevated in mouse and human muscle in disease situations characterized by unresolved inflammation, and chronic tissue remodeling, such as muscular dystrophy and cancer cachexia. This dichotomy between transient physiological activation and chronic pathological elevation is not uncommon, nor is the challenge to discern between a causative detrimental process and an attempt at rescue. In our study, ectopic Zfp697 expression in skeletal muscle over a period of 7 days led to increased collagen deposition and heightened expression of markers of inflammation and ECM remodeling, suggesting that chronically elevated Zfp697 may contribute to skeletal muscle dysfunction. As the pattern of Zfp697 expression is not restricted to skeletal muscle, it remains to be established to what extent it might contribute to the pathophysiology of different diseases.

ZFPs represent an abundant group of proteins, with diverse biological activities and mechanisms of action. ZFPs can interact with DNA, RNA, and other proteins and have as diverse cellular functions as regulation of transcription, DNA repair, signal transduction, protein degradation, among other (*16*). In skeletal muscle, ZFPs are known to regulate myoblast differentiation (i.e. KLF5) (*46*)) and fusion (i.e. KLF2/4) (*47*), protein turnover and muscle mass (i.e. Murf1, LMCD1) (*48, 49*), metabolism and exercise adaptation (i.e. YY1, KLF15) (*50, 51*), among many other processes. Zfp697 contains eleven C2H2 zinc fingers, spanning most of the protein, which led us to test if they could mediate DNA-binding. Although our data suggest that in muscle Zfp697 is not primarily a DNA-binding TF, we cannot preclude that it may have that or other activities in a cell-type or context-specific manner. However, here we show that Zfp697 is a novel RBP that preferentially binds miRNAs and distal intronic regions. Although the number of miRNA targets we identified was not high compared to other known RBPs, suggestive of a more selective protein:RNA interaction interface, Zfp697 scored very high in its specificity for miRNA binding, even when compared with the Microprocessor complex components, DROSHA and DGCR8 (*52*). Of particular interest, the top hit of the Zfp697 miRNA list was miR-206, one of the myomiRs, a set of conserved skeletal-muscle enriched miRNAs. Among these, miR-206 and miR-133b are processed from the pre-miRNA Lincmd1 (*53*). Notably, miR-206 has been previously shown to promote skeletal muscle regeneration and to delay both features of pathology in mouse models of Duchenne muscular dystrophy (DMD) and Amyotrophic Lateral Sclerosis (ALS) in mice (*28, 29*). Concordant with the idea of physiologically transient vs pathologically chronic elevation in the expression of regulators of tissue regeneration, miR-206 has been found to be increased in circulation in ALS, muscular dystrophies, and Alzheimer’s disease patients (among others), and even suggested as a disease progression biomarker (*54*). Of note, our eCLIP data also indicate an enrichment in repetitive sequences present in Zfp697-interacting RNAs (fig. S5B). Although commonly filtered out during analysis due to the inability of mapping these sequences to unique identifiers, they often represent retrotransposable elements (*55*). When actively expressed (e.g. during cellular stress) the resulting dsRNAs are potent activators of interferon responses (*56*) and have been implicated in limb regeneration (*57*), autoimmunity (*58*), and immune cell infiltration in tumors (*59*).

Our study identifies Zfp697 as a novel RBP with a prominent role is skeletal muscle inflammation and regeneration. It is important to note that Zfp697 is not skeletal-muscle specific and that the molecular and cellular processes that it regulates and important in multiple tissues and in various physiological and pathophysiological contexts. Indeed, a search of publicly available gene expression datasets, indicates that Zfp697 expression is dysregulated in situations of unresolved inflammation and fibrosis, such as kidney and heart disease (fig. S7). The biological activities, expression patterns, signal transduction and mechanisms of action we describe here, indicate that Zfp697 can operate in diverse cell types and may be a valuable therapeutic target to improve tissue regeneration and modulate tissue inflammation.

## Acknowledgments

The authors would like to acknowledge support from Science for Life Laboratory, National Genomics Infrastructure (NGI), and Uppmax for providing assistance in massive parallel sequencing and computational infrastructure.

## Funding

This work was supported by grants from The Swedish Research Council and The Novo Nordisk Foundation to J.L.R. (2016-00785 and NNF16OC0020804); the Gösta Fraenckel Foundation for Medical Research, AFM-Tèlèthon, the European Foundation for the Study of Diabetes, and the Åke Wibergs Foundation to J.C.C.; The Swedish Research Council and Olle Engqvist Foundation to J.T.L. (2019-01282/ 2022-01650 and 207-0479); The Swedish Research Council to O.E. (80358701-03); G.W.Y. is supported as an Allen Distinguished Investigator, a Paul G. Allen Frontier Group advised grant to the Paul G. Allen Family Foundation. G.W.Y. is supported by NIH Grants U41HG009889, RF1MH126719, R01NS103172, R01HG004659 and R01HG011864. P.G. received a Senior Research Fellowship (1117835) and an Investigator Grant (2017070) from the National Health and Medical Research Council of Australia. The Research Council of Lithuania (S-MIP-19-54) supported S.K. J.C.C. and I.C. were partially supported by a postdoctoral fellowship from the Swedish Society for Medical Research (SSMF). P.R.J. received a postdoctoral fellowship from the Wenner-Gren Foundations. M.L.G. was awarded a National Science Foundation Graduate Research Fellowship (DGE-2038238), a Myotonic Dystrophy Foundation Doctoral Research Fellowship, and an Association for Women in Science Scholarship. S.D. received a postdoctoral fellowship from the Swiss National Science Foundation. Q.L. was awarded an American Heart Association Predoctoral Fellowship.

## Author contributions

Conceptualization: J.C.C. and J.L.R.

Methodology: J.C.C., P.R.J., M.L.G., D.E., O.E., P.G., G.W.Y.

Investigation: J.C.C., P.R.J., M.L.G., I.C., S.D., S.M.P., K.D., Z.L., Q.L., T.V, M.B., S.K., J.T.L.

Visualization: J.C.C., P.R.J., I.G. and S.D.

Funding acquisition: J.C.C., O.E., P.G., S.K., G.W.Y and J.L.R.

Project administration: J.L.R.

Supervision: J.L.R., J.T.L., G.W.Y., H.W., S.K.

Writing – original draft: J.C.C., P.R.J. and J.L.R.

Writing – review & editing: J.L.R. and all authors

## Competing Interests

G.W.Y. is an SAB member of Jumpcode Genomics and a co-founder, member of the Board of Directors, on the SAB, equity holder, and paid consultant for Locanabio and Eclipse BioInnovations. G.W.Y. is a distinguished visiting professor at the National University of Singapore. G.W.Y.’s interests have been reviewed and approved by the University of California, San Diego in accordance with its conflict-of-interest policies.

## Data and materials availability

Unique reagents generated in this study will be made available upon reasonable request to the corresponding author with a completed Materials Transfer Agreement. All data are available in the main text or the supplementary materials. Next-generation sequencing (NGS) data generated is being deposited at GEO and will be available ahead of publication. In addition, all used datasets are made available as Data S1 to S8 (attached). Code used to analyze NGS data will be available at the RuasLab GitHub (https://github.com/ruaslab). Any additional information required to reanalyze the data reported in this paper is available from the lead contact upon request.

## Supplementary Materials

Materials and Methods

Figs. S1 to S9

References

Data S1 to S8

## References and Notes

1. L. S. Chow, R. E. Gerszten, J. M. Taylor, B. K. Pedersen, H. van Praag, S. Trappe, M. A. Febbraio, Z. S. Galis, Y. Gao, J. M. Haus, I. R. Lanza, C. J. Lavie, C.-H. Lee, A. Lucia, C. Moro, A. Pandey, J. M. Robbins, K. I. Stanford, A. E. Thackray, S. Villeda, M. J. Watt, A. Xia, J. R. Zierath, B. H. Goodpaster, M. P. Snyder, Exerkines in health, resilience and disease. Nat Rev Endocrinol. 18, 273–289 (2022).

2. R. Furrer, J. A. Hawley, C. Handschin, The molecular athlete: exercise physiology from mechanisms to medals. Physiological Reviews (2023), doi:10.1152/physrev.00017.2022.

3. S. M. Hindi, D. P. Millay, All for One and One for All: Regenerating Skeletal Muscle. Cold Spring Harb Perspect Biol. 14, a040824 (2022).

4. A. W. B. Joe, L. Yi, A. Natarajan, F. Le Grand, L. So, J. Wang, M. A. Rudnicki, F. M. V. Rossi, Muscle injury activates resident fibro/adipogenic progenitors that facilitate myogenesis. Nat Cell Biol. 12, 153–163 (2010).

5. C. Bernard, A. Zavoriti, Q. Pucelle, B. Chazaud, J. Gondin, Role of macrophages during skeletal muscle regeneration and hypertrophy-Implications for immunomodulatory strategies. Physiol Rep. 10, e15480 (2022).

6. D. Burzyn, W. Kuswanto, D. Kolodin, J. L. Shadrach, M. Cerletti, Y. Jang, E. Sefik, T. G. Tan, A. J. Wagers, C. Benoist, D. Mathis, A special population of regulatory T cells potentiates muscle repair. Cell. 155, 1282–1295 (2013).

7. M. V. Plikus, X. Wang, S. Sinha, E. Forte, S. M. Thompson, E. L. Herzog, R. R. Driskell, N. Rosenthal, J. Biernaskie, V. Horsley, Fibroblasts: Origins, definitions, and functions in health and disease. Cell. 184, 3852–3872 (2021).

8. L. R. Smith, E. R. Barton, Regulation of fibrosis in muscular dystrophy. Matrix Biol. 68–69, 602–615 (2018).

9. E. R. Morey, Spaceflight and Bone Turnover: Correlation with a New Rat Model of Weightlessness. BioScience. 29, 168–172 (1979).

10. J. L. Ruas, J. P. White, R. R. Rao, S. Kleiner, K. T. Brannan, B. C. Harrison, N. P. Greene, J. Wu, J. L. Estall, B. a Irving, I. R. Lanza, K. a Rasbach, M. Okutsu, K. S. Nair, Z. Yan, L. a Leinwand, B. M. Spiegelman, A PGC-1α isoform induced by resistance training regulates skeletal muscle hypertrophy. Cell. 151, 1319–31 (2012).

11. A. Liberzon, C. Birger, H. Thorvaldsdóttir, M. Ghandi, J. P. Mesirov, P. Tamayo, The Molecular Signatures Database Hallmark Gene Set Collection. cels. 1, 417–425 (2015).

12. S. A. Lambert, A. Jolma, L. F. Campitelli, P. K. Das, Y. Yin, M. Albu, X. Chen, J. Taipale, T. R. Hughes, M. T. Weirauch, The Human Transcription Factors. Cell. 172, 650–665 (2018).

13. S. He, T. Fu, Y. Yu, Q. Liang, L. Li, J. Liu, X. Zhang, Q. Zhou, Q. Guo, D. Xu, Y. Chen, X. Wang, Y. Chen, J. Liu, Z. Gan, Y. Liu, IRE1α regulates skeletal muscle regeneration through Myostatin mRNA decay. J Clin Invest. 131, e143737 (2021).

14. M. A. Pearen, G. E. O. Muscat, Minireview: Nuclear hormone receptor 4A signaling: implications for metabolic disease. Mol Endocrinol. 24, 1891–1903 (2010).

15. K. B. Umansky, Y. Gruenbaum-Cohen, M. Tsoory, E. Feldmesser, D. Goldenberg, O. Brenner, Y. Groner Runx1 Transcription Factor Is Required for Myoblasts Proliferation during Muscle Regeneration. PLoS Genet. 11, e1005457 (2015).

16. M. Cassandri, A. Smirnov, F. Novelli, C. Pitolli, M. Agostini, M. Malewicz, G. Melino G. Raschellà, Zinc-finger proteins in health and disease. Cell Death Discov. 3, 17071 (2017).

17. F. Sarto, D. W. Stashuk, M. V. Franchi, E. Monti, S. Zampieri, G. Valli, G. Sirago, J. Candia, L. M. Hartnell, M. Paganini, J. S. McPhee, G. De Vito, L. Ferrucci, C. Reggiani, M. V. Narici, Effects of short-term unloading and active recovery on human motor unit properties, neuromuscular junction transmission and transcriptomic profile. The Journal of Physiology. n/a (2022), doi:10.1113/JP283381.

18. X. Zhang, M. B. Trevino, M. Wang, S. J. Gardell, J. E. Ayala, X. Han, D. P. Kelly, B. H. Goodpaster, R. B. Vega, P. M. Coen, Impaired mitochondrial energetics characterize poor early recovery of muscle mass following hind limb unloading in old mice. Journals of Gerontology - Series A Biological Sciences and Medical Sciences. 73, 1313–1322 (2018).

19. P. Hettige, U. Tahir, K. C. Nishikawa, M. J. Gage, Transcriptomic profiles of muscular dystrophy with myositis (mdm) in extensor digitorum longus, psoas, and soleus muscles from mice. BMC Genomics. 23, 657 (2022).

20. T. A. Blackwell, I. Cervenka, B. Khatri, J. L. Brown, M. E. Rosa-Caldwell, D. E. Lee, R. A. Perry, L. A. Brown, W. S. Haynie, M. P. Wiggs, W. G. Bottje, T. A. Washington, B. C. Kong, J. L. Ruas, N. P. Greene, Transcriptomic analysis of the development of skeletal muscle atrophy in cancer-cachexia in tumor-bearing mice. Physiological Genomics. 50, 1071–1082 (2018).

21. N. J. Pillon, J. A. B. Smith, P. S. Alm, A. V. Chibalin, J. Alhusen, E. Arner, P. Carninci, T. Fritz, J. Otten, T. Olsson, S. van Doorslaer de ten Ryen, L. Deldicque, K. Caidahl, H. Wallberg-Henriksson, A. Krook, J. R. Zierath, Distinctive exercise-induced inflammatory response and exerkine induction in skeletal muscle of people with type 2 diabetes. Science Advances. 8, eabo3192 (2022).

22. M. J. Petrany, C. O. Swoboda, C. Sun, K. Chetal, X. Chen, M. T. Weirauch, N. Salomonis, D. P. Millay, Single-nucleus RNA-seq identifies transcriptional heterogeneity in multinucleated skeletal myofibers. Nat Commun. 11, 6374 (2020).

23. A. J. De Micheli, E. J. Laurilliard, C. L. Heinke, H. Ravichandran, P. Fraczek, S. Soueid-Baumgarten, I. De Vlaminck, O. Elemento, B. D. Cosgrove, Single-Cell Analysis of the Muscle Stem Cell Hierarchy Identifies Heterotypic Communication Signals Involved in Skeletal Muscle Regeneration. Cell Rep. 30, 3583-3595.e5 (2020).

24. V. Gotea, I. Ovcharenko, DiRE: identifying distant regulatory elements of co-expressed genes. Nucleic Acids Research. 36, W133–W139 (2008).

25. M. Cheng, M.-H. Nguyen, G. Fantuzzi, T. J. Koh, Endogenous interferon-gamma is required for efficient skeletal muscle regeneration. Am J Physiol Cell Physiol. 294, C1183–1191 (2008).

26. C. Zhang, N. Cheng, B. Qiao, F. Zhang, J. Wu, C. Liu, Y. Li, J. Du, Age-related decline of interferon-gamma responses in macrophage impairs satellite cell proliferation and regeneration. J Cachexia Sarcopenia Muscle. 11, 1291–1305 (2020).

27. E. L. Van Nostrand, G. A. Pratt, A. A. Shishkin, C. Gelboin-Burkhart, M. Y. Fang, B. Sundararaman, S. M. Blue, T. B. Nguyen, C. Surka, K. Elkins, R. Stanton, F. Rigo, M. Guttman, G. W. Yeo, Robust transcriptome-wide discovery of RNA-binding protein binding sites with enhanced CLIP (eCLIP). Nat Methods. 13, 508–514 (2016).

28. N. Liu, A. H. Williams, J. M. Maxeiner, S. Bezprozvannaya, J. M. Shelton, J. A. Richardson, R. Bassel-Duby, E. N. Olson, microRNA-206 promotes skeletal muscle regeneration and delays progression of Duchenne muscular dystrophy in mice. J Clin Invest. 122, 2054–2065 (2012).

29. A. H. Williams, G. Valdez, V. Moresi, X. Qi, J. McAnally, J. L. Elliott, R. Bassel-Duby, J. R. Sanes, E. N. Olson, MicroRNA-206 delays ALS progression and promotes regeneration of neuromuscular synapses in mice. Science. 326, 1549–1554 (2009).

30. D. P. Bartel, MicroRNAs: genomics, biogenesis, mechanism, and function. Cell. 116, 281– 297 (2004).

31. S. Li, M. P. Czubryt, J. McAnally, R. Bassel-Duby, J. A. Richardson, F. F. Wiebel, A. Nordheim, E. N. Olson, Requirement for serum response factor for skeletal muscle growth and maturation revealed by tissue-specific gene deletion in mice. Proc. Natl. Acad. Sci. U.S.A. 102, 1082–1087 (2005).

32. A. M. Newman, C. B. Steen, C. L. Liu, A. J. Gentles, A. A. Chaudhuri, F. Scherer, M. S. Khodadoust, M. S. Esfahani, B. A. Luca, D. Steiner, M. Diehn, A. A. Alizadeh, Determining cell type abundance and expression from bulk tissues with digital cytometry. Nat Biotechnol. 37, 773–782 (2019).

33. X. J. Musacchia, J. M. Steffen, D. R. Deavers, Rat hindlimb muscle responses to suspension hypokinesia/hypodynamia. Aviat Space Environ Med. 54, 1015–1020 (1983).

34. F. Chemello, Z. Wang, H. Li, J. R. McAnally, N. Liu, R. Bassel-Duby, E. N. Olson, Degenerative and regenerative pathways underlying Duchenne muscular dystrophy revealed by single-nucleus RNA sequencing. Proceedings of the National Academy of Sciences. 117, 29691–29701 (2020).

35. J. G. Tidball, Regulation of muscle growth and regeneration by the immune system. Nat Rev Immunol. 17, 165–178 (2017).

36. P. J. Ferrara, P. T. Reidy, J. J. Petrocelli, E. M. Yee, D. K. Fix, Z. S. Mahmassani, J. A. Montgomery, A. I. McKenzie, N. M. M. P. de Hart, M. J. Drummond, Global deletion of CCL2 has adverse impacts on recovery of skeletal muscle fiber size and function and is muscle specific. Journal of Applied Physiology. 134, 923–932 (2023).

37. H. Koike, I. Manabe, Y. Oishi, Mechanisms of cooperative cell-cell interactions in skeletal muscle regeneration. Inflamm Regen. 42, 48 (2022).

38. H. Lu, D. Huang, R. M. Ransohoff, L. Zhou, Acute skeletal muscle injury: CCL2 expression by both monocytes and injured muscle is required for repair. The FASEB Journal. 25, 3344– 3355 (2011).

39. C. Zhang, C. Wang, Y. Li, T. Miwa, C. Liu, W. Cui, W.-C. Song, J. Du, Complement C3a signaling facilitates skeletal muscle regeneration by regulating monocyte function and trafficking. Nat Commun. 8, 2078 (2017).

40. S. Zhuang, A. Russell, Y. Guo, Y. Xu, W. Xiao, IFN-*γ* blockade after genetic inhibition of PD-1 aggravates skeletal muscle damage and impairs skeletal muscle regeneration. Cell Mol Biol Lett. 28, 27 (2023).

41. S. Mahurkar, M. Moldovan, V. Suppiah, M. Sorosina, F. Clarelli, G. Liberatore, S. Malhotra, X. Montalban, A. Antigüedad, M. Krupa, V. G. Jokubaitis, F. C. McKay, P. N. Gatt, M. J. Fabis-Pedrini, V. Martinelli, G. Comi, J. Lechner-Scott, A. G. Kermode, M. Slee, B. V. Taylor, K. Vandenbroeck, M. Comabella, F. M. Boneschi, Australian and New Zealand Multiple Sclerosis Genetics Consortium (ANZgene), C. King, Response to interferon-beta treatment in multiple sclerosis patients: a genome-wide association study. Pharmacogenomics J. 17, 312–318 (2017).

42. N. A. Dumont, C. F. Bentzinger, M.-C. Sincennes, M. A. Rudnicki, Satellite Cells and Skeletal Muscle Regeneration. Compr Physiol. 5, 1027–1059 (2015).

43. M. N. Wosczyna, C. T. Konishi, E. E. P. Carbajal, T. T. Wang, R. A. Walsh, Q. Gan, M. W. Wagner, T. A. Rando, Mesenchymal Stromal Cells Are Required for Regeneration and Homeostatic Maintenance of Skeletal Muscle. Cell Reports. 27, 2029-2035.e5 (2019).

44. S. J. Forbes, N. Rosenthal, Preparing the ground for tissue regeneration: from mechanism to therapy. Nat Med. 20, 857–869 (2014).

45. C. J. Mann, E. Perdiguero, Y. Kharraz, S. Aguilar, P. Pessina, A. L. Serrano, P. Mu ñ oz-Cánoves, Aberrant repair and fibrosis development in skeletal muscle. Skelet Muscle. 1, 21 (2011).

46. S. Hayashi, I. Manabe, Y. Suzuki, F. Relaix, Y. Oishi, Klf5 regulates muscle differentiation by directly targeting muscle-specific genes in cooperation with MyoD in mice. Elife. 5, e17462 (2016).

47. K. Sunadome, T. Yamamoto, M. Ebisuya, K. Kondoh, A. Sehara-Fujisawa, E. Nishida, ERK5 regulates muscle cell fusion through Klf transcription factors. Dev Cell. 20, 192–205 (2011).

48. S. C. Bodine, E. Latres, S. Baumhueter, V. K.-M. Lai, L. Nunez, B. A. Clarke, W. T. Poueymirou, F. J. Panaro, E. Na, K. Dharmarajan, Z.-Q. Pan, D. M. Valenzuela, T. M. DeChiara, T. N. Stitt, G. D. Yancopoulos, D. J. Glass, Identification of Ubiquitin Ligases Required for Skeletal Muscle Atrophy. Science. 294, 1704–1708 (2001).

49. D. M. S. Ferreira, A. J. Cheng, L. Z. Agudelo, I. Cervenka, T. Chaillou, J. C. Correia, M. Porsmyr-Palmertz, M. Izadi, A. Hansson, V. Mart Í nez-Redondo, P. Valente-Silva, A. T. Pettersson-Klein, J. L. Estall, M. M. Robinson, K. S. Nair, J. T. Lanner, J. L. Ruas, LIM and cysteine-rich domains 1 (LMCD1) regulates skeletal muscle hypertrophy, calcium handling, and force. Skeletal Muscle. 9, 26 (2019).

50. J. T. Cunningham, J. T. Rodgers, D. H. Arlow, F. Vazquez, V. K. Mootha, P. Puigserver, mTOR controls mitochondrial oxidative function through a YY1–PGC-1α transcriptional complex. Nature. 450, 736–740 (2007).

51. S. M. Haldar, D. Jeyaraj, P. Anand, H. Zhu, Y. Lu, D. a Prosdocimo, B. Eapen, D. Kawanami, M. Okutsu, L. Brotto, H. Fujioka, J. Kerner, M. G. Rosca, O. P. McGuinness, R. J. Snow, A. P. Russell, A. N. Gerber, X. Bai, Z. Yan, T. M. Nosek, M. Brotto, C. L. Hoppel, M. K. Jain, Kruppel-like factor 15 regulates skeletal muscle lipid flux and exercise adaptation. Proceedings of the National Academy of Sciences of the United States of America. 109, 6739–44 (2012).

52. M. Ha, V. N. Kim, Regulation of microRNA biogenesis. Nat Rev Mol Cell Biol. 15, 509–524 (2014).

53. M. Cesana, D. Cacchiarelli, I. Legnini, T. Santini, O. Sthandier, M. Chinappi, A. Tramontano, I. Bozzoni, A Long Noncoding RNA Controls Muscle Differentiation by Functioning as a Competing Endogenous RNA. Cell. 147, 358–369 (2011).

54. S. Khalilian, S. Z. Hosseini Imani, S. Ghafouri-Fard, Emerging roles and mechanisms of miR-206 in human disorders: a comprehensive review. Cancer Cell Int. 22, 412 (2022).

55. E. L. Van Nostrand, G. A. Pratt, B. A. Yee, E. C. Wheeler, S. M. Blue, J. Mueller, S. S. Park, K. E. Garcia, C. Gelboin-Burkhart, T. B. Nguyen, I. Rabano, R. Stanton, B. Sundararaman, R. Wang, X.-D. Fu, B. R. Graveley, G. W. Yeo, Principles of RNA processing from analysis of enhanced CLIP maps for 150 RNA binding proteins. Genome Biol. 21, 90 (2020).

56. A. M. Herzner, Z. Khan, E. L. van Nostrand, S. Chan, T. Cuellar, R. Chen, X. Pechuan-Jorge, L. Komuves, M. Solon, Z. Modrusan, B. Haley, G. W. Yeo, T. W. Behrens, M. L. Albert, ADAR and hnRNPC deficiency synergize in activating endogenous dsRNA-induced type I IFN responses. Journal of Experimental Medicine. 218 (2021), doi:10.1084/jem.20201833.

57. W. Zhu, D. Kuo, J. Nathanson, A. Satoh, G. M. Pao, G. W. Yeo, S. V. Bryant, S. R. Voss, D. M. Gardiner, T. Hunter, Retrotransposon long interspersed nucleotide element-1 (LINE-1) is activated during salamander limb regeneration. Dev Growth Differ. 54, 673–685 (2012).

58. M. J. Heinrich, C. A. Purcell, A. J. Pruijssers, Y. Zhao, C. F. Spurlock, S. Sriram, K. M. Ogden, T. S. Dermody, M. B. Scholz, P. S. Crooke, J. Karijolich, T. M. Aune, Endogenous double-stranded Alu RNA elements stimulate IFN-responses in relapsing remitting multiple sclerosis. J Autoimmun. 100, 40–51 (2019).

59. Y. Kong, C. M. Rose, A. A. Cass, A. G. Williams, M. Darwish, S. Lianoglou, P. M. Haverty, A.-J. Tong, C. Blanchette, M. L. Albert, I. Mellman, R. Bourgon, J. Greally, S. Jhunjhunwala, H. Chen-Harris, Transposable element expression in tumors is associated with immune infiltration and increased antigenicity. Nat Commun. 10, 5228 (2019).

