## Supplementary Methods and Figures for "Zfp697 is an RNA-binding protein that regulates skeletal muscle inflammation and regeneration"

### Materials and Methods

#### Animal experiments

Mice were housed in groups of 3-5 per cage on a 12-hour light/dark cycle, at approximately 22° C (GM500 cages, Tecniplast). Animals had access to standard rodent chow diet (Special Diets Services CRM-P) and water *ad libitum*. Only healthy sex- and age-matched mice were used in experiments, and their ages are indicated in the respective Method Details section. The Zfp697-mKO mice and their flox littermates were generated in-house, while C57BL/6N, C57BL/6J, and CB-17/lcr-Prkdcscid/Rj (SCID) mice were purchased from Janvier Labs and acclimated to the animal facility before use. All tissues were collected, immediately frozen in liquid nitrogen, and stored at -80°C until analysis. All experiments were approved by the regional animal ethics committee of Northern Stockholm, Sweden.

#### Hindlimb unloading and reloading

This protocol was modified from previous studies (*1*). 10 to 12-week-old male C57BL/6J mice or 8-week-old female Zfp697-mKO mice and flox littermates were assigned to three main groups: control, hindlimb unloading or hindlimb reloading. For the hindlimb unloading phase, mice were slightly suspended by tail for the specified number of days, up to 10 days. Briefly, mice were placed inside a restrainer and, using hypoallergenic medical tape, a nylon line was attached in a helical pattern around the base of the tail. The line was connected to a small swivel keychain with metal rings that were attached to a rod running the length of the cage (GR900, Tecniplast). To prevent their hind legs from carrying any weight, the hindlimbs were kept just off the ground with the mouse's body at an approximately 30° angle. Mice were able to move freely on the y-axis, rotate 360° using their forelegs, and had access to one side of the cage with food and water *ad libitum*. Mice were housed in pairs during the unloading phase. The reloading phase consisted of 10 days of hindlimb unloading, after which the animals were returned to normal ambulation in conventional cages for the specified number of days. Control mice were kept in conventional cages throughout the entire protocol with food and water *ad libitum*.

#### Chemically induced muscle injury

To induce muscle injury, mice were intramuscularly injected with either barium chloride ($BaCl_2$; 1.2% in saline) or cardiotoxin (10 μM). Prior to the procedure, mice were given a subcutaneous injection of 0.05 - 0.1 mg/kg buprenorphine and placed under isoflurane anesthesia. For experiments using $BaCl_2$, right gastrocnemius muscle of 8-week-old male C57BL/6J mice were injected with 50 μL of $BaCl_2$, while the contralateral gastrocnemius was injected with an equivalent volume of control solution (saline). For experiments using cardiotoxin, 10-week-old female Zfp697-mKO and floxed littermates were injected with 30 μL of cardiotoxin into the right gastrocnemius muscle and 20 μL into the right tibialis anterior muscle. Mice were monitored and given postoperative pain relief for 1-2 days. The injected muscles were collected at the specified time points.

#### Clenbuterol injections

8-week-old male C57BL/6J mice were intraperitoneally injected with 2 mg/kg body weight of clenbuterol (Sigma-Aldrich) or equivalent volume of PBS. Gastrocnemius muscles were harvested 16 hours post-injection.

**LPS injections**
8-week-old male C57BL/6J mice were intraperitoneally injected with 1 μg/g body weight of LPS (Ultrapure lipopolysaccharide from E-Coli K12, Invivogen) or equivalent volume of vehicle. Soleus muscles were harvested 3 hours post-injection.

**Primary myoblasts isolation**
Mouse primary myoblasts were isolated, cultured and differentiated according to previously described methods (*2*). Muscles of 2-week-old C57BL/6J mice were harvested from the fore- and hindlimbs, minced, and incubated with 2.4 U/ml dispase (Grade II, Roche), 1% collagenase B (Roche), and 2.5 mM $CaCl_2$ for 25 minutes at 37°C. The cell suspension was homogenized by pipetting, filtered through a 70 μm cell strainer, centrifuged, resuspended in Ham's F-10 Nutrient Mix (Gibco) supplemented with 20% fetal bovine serum (Sigma-Aldrich), 2.5 ng/mL basic fibroblast growth factor (Thermo-Fisher Scientific), 2.5 mg/ml fungizone (Thermo-Fisher Scientific), 5 mg/ml plasmocin (Invivogen) and 100 U/ml penicillin-streptomycin (Gibco), and plated in regular tissue culture dishes to remove contaminant fibroblasts. After 20 minutes, cells were transferred to collagen-coated tissue culture dishes. Pre-plating in regular culture dishes was performed at every passage until there was no fibroblast contamination. Cells were then maintained and expanded at low confluence in a 1:1 mixture of DMEM (Gibco) and Ham's F10 Nutrient Mix supplemented with 20% fetal bovine serum, 2.5 ng/ml basic fibroblast growth factor and 100 U/ml penicillin-streptomycin. To differentiate the myoblasts into myotubes, cells were seeded in growth medium at high confluence and switched to differentiation medium (DMEM with 5% horse serum and 100 U/ml penicillin-streptomycin) 16-24 hours later. Full differentiation of the myotubes was typically achieved 2-3 days after induction and was confirmed using light microscopy. Myoblasts were maintained, expanded, and differentiated in collagen-coated tissue culture dishes at 37°C and 5% CO2.

**Generation of a muscle-specific Zfp697 knockout mouse model**
Floxed mice for the conditional deletion of the *Zfp697* gene were generated by Biocytogen (USA), using CRISPR/Cas9-mediated homology-directed repair. Briefly, loxP sites were inserted in the intronic region upstream of exon 3 and within the 3´UTR region of exon 3. To minimize the possibility of disrupting *Zfp697* expression, both loxP sites were inserted into regions of the gene with low sequence conservation between species. Genomic DNA was extracted from F1 pups and southern blot was used to screen out animals with random insertions. Muscle-specific Zfp697 knockouts (Zfp697-mKO) were generated by crossing *Zfp697* floxed mice with animals expressing Cre recombinase under the control of the 1.5-kb *Myog* promoter and the 1-kb *Mef2c* enhancer (Myo-Cre; a generous gift from R. Bassel-Duby and E.N. Olson, University of Texas Southwestern Medical Center, USA) (*3*). Primers for genotyping *Zfp697* floxed allele were the following: 5´loxP forward, 5´-TCTGGGACCTGATGAGTCCTACTGG-3´; 5´loxP reverse, 5´-ACGGAGGATAGCCCTTTCTTCCTCA.

**Human high-intensity interval training**
For analysis of *ZNF697* gene expression in human muscle after high-intensity exercise, we used vastus lateralis muscle biopsies from a previously published study (*4*). Recreationally active subjects and elite endurance athletes completed a single session of high-intensity interval training (HIIT) on a cycle ergometer, consisted of three to six 30-second cycling intervals at 0.7 Nm per kg of body weight, with a 4-minute rest period in between each interval. Muscle samples were taken before and 24 hours after the exercise session and used for mRNA analyses.

**Human unloading and reloading**

For analysis of *ZNF697* gene expression in human muscle after limb unloading muscle and recovery, we used vastus lateralis muscle biopsies from a subset of subjects of a previously published study (*5*). Briefly, recreationally active young and old male subjects underwent 3 days of unloading of the dominant leg by wearing a 10 cm extended-sole shoe on the contralateral leg and allowed to move (< 2000 steps per day) with the aid of forearm crutches. After the unloading period, subjects returned to normal ambulatory activity and were enrolled in a 3-week resistance training regimen as described before (*5*). Biopsies were collected from the unloaded leg at baseline, after the unloading period and at the end of the 3 weeks of resistance training. This study was approved by the Regional Biomedical Research Ethics Committee (no. BE-2-47).

**Grip strength test**

Mouse forelimb grip strength was determined using a grip strength meter (Bioseb). Mice were held by their tails and allowed to grasp the bar of the device with their front paws. They were then gently pulled away from the bar until they let go, and the force applied was recorded. This process was repeated three times, for up to 4 consecutive days, with the mice being weighed for normalization prior to the first measurement.

**Exercise protocols**

Treadmill running to exhaustion test

Mice were acclimated to the treadmill (Columbus Instruments, Columbus, OH) for 4 consecutive days before the experiment. On the fifth day, the mice began running at a speed of 6 m/min on a 10° upward incline, with the speed being increased by 3 m/min every 3 minutes until the point of exhaustion. Exhaustion was identified as the moment when the mice refused to continue running even with gentle encouragement. Maximal speed, total distance and time were recorded. The exercise performance of Zfp697-mKO mice and flox littermates was evaluated using 14-week-old males.

Acute treadmill running

Two days after a run to exhaustion test, 8-week-old male C57BL/6J mice were submitted to an acute bout of treadmill running performed at 60% of the maximal speed achieved during the test, with a 10° upward incline, for 60 minutes. All exercise bouts were performed at the same time of the day (ZT 18), and animals were euthanized and had their quadriceps muscles harvested immediately, 3 hours, 6 hours, 12 hours or 24 hours after running. Rest control mice were subjected to the same procedures except the acute bout of exercise, and euthanized at different times of the day and in parallel with the exercised mice.

Downhill treadmill running

To induce eccentric muscle damage, ~9-week-old male mice were acclimated to the treadmill as described above. One day after the acclimatization period, animals were submitted to an acute bout of downhill running, consisted of running for 5 min at 6 m/min (warm-up), followed by 55 minutes at 17 m/min, on a 15° downward slope.

**Body composition measurement**

Mice were anaesthetized using isoflurane and their body composition was examined using dual-energy X-ray absorptiometry (DEXA) on a Lunar PIXImus densitometer (GE Medical Systems).

### Gait Analysis

Gait analysis was performed on Zfp697-mKO and flox littermate controls that received a unilateral intramuscular injection of cardiotoxin. One day before and 1, 2, 3 and 5 days after the injection, gait analysis was conducted using the DigiGait treadmill and software (Mouse Specific Inc.), which incorporates Ventral Plane Imaging Technology to create digital representations of paw prints from the anima while it moves on a motorized treadmill. Briefly, each mouse was placed on a transparent treadmill belt and the speed was set to 15 cm/s (brisk walk speed). Ventral view of the animals as they walk was recorded and a segment of 5 seconds of uninterrupted walking was used. Each video was then processed through the DigiGait Analysis software, as described previously (*6*), and parameters for stride length, stance (duration of paw contact with the ground), swing time (duration of no paw contact with the ground), and propulsion (duration of maximum paw contact to beginning of swing phase) were extracted.

### Analysis of gene expression

Total RNA was extracted from cell cultures and frozen tissue using TRI reagent (Sigma-Aldrich) as per the manufacturer's instructions. One μg of RNA was treated with Amplification Grade DNAse I (Thermo Scientific) and 500 ng of DNase-treated RNA was used to prepare cDNA using the High-Capacity cDNA Reverse Transcription Kit (Applied Biosystems). Quantitative real-time PCR was conducted on ViiA7 and QuantStudio6 real-time PCR systems (Applied Biosystems) using SYBR Green PCR Master Mix (Applied Biosystems). Gene expression was calculated by the delta-delta Ct method and normalized to the expression of TATA-binding protein (Tbp).

### Global gene expression analysis by RNA-sequencing

Total RNA was extracted from primary myotubes and gastrocnemius muscles of mice using TRI reagent (Sigma-Aldrich) as per the manufacturer's instructions. RNA was purified and treated with DNase using NucleoSpin RNA II columns (Machery Nagel) to remove any remaining DNA contaminants. RNA integrity was verified using an Agilent Bioanalyzer, with all samples showing RNA integrity number (RIN) greater than 8. Stranded mRNA Library Preparation Kit was used with a polyA-enrichment strategy to prepare the sequencing libraries. RNA-sequencing for the initial hindlimb unloading and reloading experiment was performed at the SciLifeLab (Stockholm, Sweden) using the Illumina HiSeq2000, generating 100-bp paired-end reads. RNA-sequencing for myotubes overexpressing Zfp697 was performed at GATC Biotech (Konstanz, Germany) using the Illumina HiSeq2500, generating 50-bp single-end reads. RNA-sequencing for hindlimb unloading and reloding using Zfp697-mKO and flox littermates was performed by the University of Chicago Genomics Facility (Chicago, USA) using the Illumina NovaSeq, generating 100-bp paired-end reads. Quality control of raw reads was performed using the FastQC tool kit (Babraham Bioinformatics). The reads were then aligned to the GRCm38.p6 mouse genome using STAR (*7*). Feature count was performed using featureCounts (*8*), and differential expression analyses were generated in R using the DESeq2 package (*9*) with adaptive log-fold change shrinkage estimator from the *ashr* package (*10*). Pathway analysis was carried out by Gene Set Enrichment Analysis (GSEA) with pre-ranked list of genes ($-\log_{10}$(p-value) * sign($\log_2$Fold-Change) (*11*) and hallmark pathways gene sets (*12*), or AmiGO (*13*) for overrepresentation analysis. The data files are being deposited in NCBI GEO.

### Digital cytometry with CIBERSORTx

Deconvolution of Zfp697-mKO hindlimb unloading/reloading bulk RNA sequencing data was performed using CIBERSORTx (*14*) (https://cibersortx.stanford.edu/) with single cell RNA sequencing data of regenerating muscle (GSE143437) (*15*) providing cell type expression signatures. The signature matrix was generated using normalized counts from single cell sequencing and with quantile normalization disabled (default parameters). Using this signature matrix, cell fractions were imputed from normalized counts from bulk RNA sequencing without batch correction and with quantile normalization disabled, using 100 permutations.

**Chromatin Immunoprecipitation**

Fully differentiated mouse primary myotubes were transduced with adenovirus expressing Flag-tagged Zfp697. Cells were crosslinked 36-40 hours post-infection by adding 1% formaldehyde directly into the media for 12 minutes at room temperature with gentle shaking. The crosslinking reaction was quenched by adding glycine to a final concentration of 125 mM. Cells were washed and collected with ice-cold PBS, and centrifuged at 850g for 5 minutes at room temperature. The pellet was resuspended in 1 mL of ChIP Buffer I (0.25% Triton X-100, 10 mM EDTA, 0.5 mM EGTA, 10 mM HEPES pH 6.5) and centrifuged for 5 minutes at 850g at 4°C. The pellet was then resuspended in 1 mL of ChIP Buffer II (200 mM NaCl, 1 mM EDTA, 0.5 mM EGTA, 10 mM HEPES, pH 7.5) and centrifuged again at 850g for 5 minutes at 4°C. The pellet was resuspended in 0.5 mL of ChIP Lysis Buffer (1% SDS, 10 mM EDTA, 50 mM Tris-HCl pH 8.0) supplemented with protease inhibitor cocktail (Roche, EDTA-free) and homogenized. The mixture was then incubated on ice for 30 minutes. The chromatin was then submitted to 5 cycles of sonication (Bioruptor Plus, Diagenode; 30 seconds ON, 30 seconds OFF) at low intensity in cold water. The chromatin was centrifuged at 15,000g for 10 minutes at 4°C, and the supernatant was transferred to a clear tube, and diluted 1:5 in ChIP dilution buffer (1% Triton X-100, 2 mM EDTA, 150 mM NaCl, 20 mM Tris-HCl pH 8.0) supplemented with protease inhibitor cocktail. To prepare the protein G Sepharose beads, six washes with ChIP dilution buffer were performed, and then blocked with 2 mg/mL BSA in ChIP dilution buffer for 30 minutes at 4°C with top over end rotation. The beads were centrifuged at 850g for 30 seconds and resuspended with ChIP dilution buffer to make a 50% slurry. For the chromatin immunoprecipitation, 27.5 μg of crosslinked chromatin from fully differentiated mouse primary myotubes expressing Flag-tagged Zfp697 was added to 2 μg of herring sperm DNA, 6 μg normal rabbit IgG (#2729, Cell Signaling) and 50 μL of the prepared bead slurry. The volume was adjusted to 1.1 mL with ChIP dilution buffer, and the mixture was incubated for 2 hours at 4°C with top over end rotation. The supernatant was collected after centrifugation, and 10% of the chromatin was removed as inputs and stored at -20°C. The remaining chromatin (25 μg) was incubated overnight at 4°C with top over end rotation with either 2 μg of anti-FLAG (F3165, Sigma) or normal rabbit IgG antibodies. After this step, 50 μL of BSA-blocked bead slurry was added and incubated at room temperature for 1 hour. The beads were then washed sequentially with TSE I (0.1% SDS, 1% Triton X-100, 2 mM EDTA, 20 mM Tris-HCl pH 8.0, 150 mM NaCl), TSE II (0.1% SDS, 1% Triton X-100, 2 mM EDTA, 20 mM Tris.HCl, pH8.1, 500 mM NaCl), ChIP buffer III (0.25 M LiCl, 1% NP-40, 1% deoxycholate, 1 mM EDTA, 10 mM Tris-HC, pH 8.0) and three times with TE buffer (10 mM Tris-HCl pH 8.0, 1 mM EDTA) at room temperature, with top over end rotation. Elution was performed three times with 100 μL of ChIP elution buffer (1% SDS, 0.1 M $NaHCO_3$) at room temperature with gentle agitation. The eluates were combined (total volume of 300 μL) and 12 μL of 5M NaCl was added. The inputs were processed in parallel. The mixture was incubated overnight at 65°C. Next, 3 μL of RNase was added, and the sample was incubated for 1 hour at 37°C, followed by the addition of 6 μL of Proteinase K and incubation for 4 hours at 55°C. DNA was extracted with phenol/chloroform/isoamyl, followed by ethanol precipitation with glycogen at a final concentration of 5 µg/mL. The DNA was resuspended in 50 µL of nuclease-free H2O and used for library preparation. The libraries for ChIP-seq were generated using the Takara ThruPLEX DNA-Seq Kit (Takara, R400736). Subsequently, single-ended sequencing was conducted on an Illumina NextSeq 550 (Illumina, 75SE reads) by the BEA Core Facility (Karolinska Institutet, Sweden). The ChIP-seq library was prepared using Takara ThruPLEX DNA-Seq Kit (Takara, R400736). Sequencing was performed in the Illumina NextSeq 550 (Illumina, 75SE reads) by the BEA Core Facility) with single-ended reads. Reads were aligned to the mm10 version of the mouse genome using STAR (*7*) and duplicated reads were removed with MarkDuplicates. Next, we used MACS3 callpeak for peak calling (q-value < 0.05 and fold change ≥ 1.5) (*16*), and bedtools to generate coverage tracks (*17*).

**Enhanced crosslinking and immunoprecipitation (eCLIP)**
eCLIP was performed as previously described (*18*). Briefly, primary mouse myotubes were UV-cross-linked (400 mJ cm-2, 254 nm) and lysed. Lysates were sonicated and treated with RNase I to fragment RNA. Two percent of each lysate sample was reserved for preparation of a parallel size-matched input library. The remaining lysates were immunoprecipitated using anti-FLAG antibody (Sigma Aldrich, F1804). Bound RNA fragments were dephosphorylated and 3'-end ligated to an RNA adaptor. Reverse transcription was performed with AffinityScript (Agilent), and cDNAs were 5'-end ligated with a DNA adaptor. cDNA yields were quantified by qPCR, and libraries were amplified using Q5 PCR mix (NEB). Libraries were sequenced on the Illumina NovaSeq6000 to a depth of at least 25×10^6 reads per library. Sequencing reads were processed as described previously (*18*). Briefly, reads were adapter-trimmed and mapped to mouse-specific repetitive elements from RepBase (version 18.05) by STAR (v2.7.6a). Reads aligning to repetitive genomic regions were removed, and remaining reads were then mapped to mm10. Peaks were called on the usable reads by CLIPper (*19*) and normalized to the size-matched input. Reproducible enriched peaks were generated by irreproducible discovery rate (IDR) analysis and assigned to gene regions annotated in GENCODE (v15). Peaks were deemed significant at 8-fold enrichment ($\log_2 = 3$), p-value < 0.001 ($-\log_{10} = 3$), and IDR < 0.01.

**Immunohistochemistry**
Muscles were excised and mounted on a cork support with tragacanth gum, frozen in liquid nitrogen-cooled 2-methylbutane, and 10 µm thick cross-sections were cut using a cryostat. The sections were transferred to Superfrost Plus Adhesion slides and processed for staining, with four non-consecutive sections used per sample. For immunostaining, the sections were equilibrated, fixed at -20°C with acetone for 10 minutes, followed by two washes with TBS 0.05% Triton X-100, incubated with blocking solution (TBS 0.05% Triton X-100 with 1% BSA and 10% normal serum) for 1 hour at room temperature and primary antibodies overnight at 4°C. For visualization of extracellular matrix, sections were stained with rabbit polyclonal anti-fibronectin (1:100, ab2413, Abcam) or anti-collagen I (1:100, ab34710, Abcam) antibodies. After washing, sections were incubated with secondary antibodies (1:100) for 1 hour at room temperature, washed again and mounted with ProLong™ Diamond Antifade Mountant with DAPI (Invitrogen). Sections were imaged using a Zeiss Axio Imager M2 microscope and quantifications were performed using Photoshop.
For analysis of KI67-positive cells, 10 µm thick cross-sections were cut using a cryostat and transferred to Superfrost Plus Adhesion slides. Sections were fixed with 2% PFA for 5 min, permeabilized with 0.1% Triton X-100 for 10 minutes at RT and blocked with 10% normal goat

serum (NGS) for 1h at RT. Sections were then incubated with rat anti-KI67 antibody (Thermo Fisher Scientific SOLA15, 1:100) in 10% NGS overnight at 4°C. Sections were washed with PBS and incubated with Alexa Fluor 594-conjugated goat anti-rat antibody (1:400) 1h at RT, washed wit PBS and subsequently incubated with Hoechst (1:1000) for 2 minutes at RT. Sections were finally washed and mounted with Vectashield Mounting Medium (Vector Laboratories). Imaging was performed in a Zeiss Axio Imager and quantified in ImageJ.

**Ex vivo force measurements**

Skeletal muscle force-producing capacity, fatigue resistance and recovery capacity were measured for EDL and soleus muscle from 8-9-week-old female Zfp697-mKO and age-matched floxed littermates. Mice were placed under isoflurane anesthesia, cervically dislocated, soleus and EDL muscles were immediately removed, immersed in Tyrode solution (121 mM NaCl, 5 mM KCl, 1.8 mM $CaCl_2$, 0.4 mM $NaH_2PO_4$, 0.5 mM $MgCl_2$, 24 mM $NaHCO_3$, 0.1 mM EDTA, and 5.5 mM glucose) and placed in a stimulation chamber with proximal and distal tendons attached to hooks. The chamber was maintained at 31°C using a circulatory water bath, and the Tyrode solution's pH was kept at 7.4 by continuous superfusion with carbogen (95% O2, 5% CO2). The muscles were set to optimal length, and force-frequency relationships were determined by stimulating the muscles at different frequencies (1 Hz to 120 Hz for soleus and 1 Hz to 150 Hz for EDL). Muscle-specific force ($kN/m^2$) was calculated using the muscle CSA, which was obtained by dividing muscle mass by the product of muscle length and density (1.06 g/cm3). Fatigue resistance was evaluated by measuring tetanic force at 70 Hz stimulation with 600 ms duration and 2-second intervals for 100 contractions in the soleus or at 100 Hz with 300 ms duration and 2-second intervals for 50 contractions in the EDL. After the fatigue protocol, the muscles were allowed to recover for 10 minutes, and tetanic force was measured at 1, 2, 5, and 10 minutes to determine the recovery capacity.

**Generation of recombinant adenoviruses**

Full-length *Zfp697* mouse RNA was amplified by PCR and cloned into the NotI/XbaI sites of a pcDNA3.1 plasmid (Invitrogen), in-frame with a FLAG tag inserted between the BamHI and NotI sites. The FLAG-Zfp697 cDNA was then subcloned into the pAdTrack-CMV vector (Stratagene) and the AdEasy adenoviral vector system (Stratagene) was used to generate adenovirus expressing FLAG-Zfp697, which was then amplified in Ad-293 cells and purified through CsCl gradient centrifugation, as previously described (*20*). Adenovirus expressing only GFP was used as control. For loss-of-function experiments, adenoviruses expressing Zfp697-specific shRNA (#shADV-276960; target sequence: ACCCACAGGATGATGATCTAA), or scrambled shRNA control (#1122), were acquired from VectorBiolabs. Adenoviruses used in all experiments express GFP from an independent CMV promoter.

**Adenovirus-mediated gene delivery**

Fully differentiated myotubes were transduced overnight with an adenovirus at an MOI of 100 (calculated based on number of myoblasts seeded). For intramuscular delivery of adenovirus, 14-days-old SCID mice were placed under anesthesia and 20 uL of adenovirus solution in PBS were injected in each leg ($2x10^{10}$ infectious particles). The needle was inserted into the gastrocnemius from the lower end of the muscle and parallel to the fiber orientation to minimize damage. Seven days post-injection, mice were euthanized, and tissues were collected for further analyses.

**RNA-binding assays**

The full-length Zfp697 CDS and the sequence encoding amino acids 1-192 were amplified by PCR and cloned into pGEX-4T3, in frame with GST. BL21 bacteria were transformed with the constructs (as well as an empty construct expressing GST alone) and used to express and purify GST-fused Zfp67. Briefly, BL21 bacteria transformed with each construct were inoculated into 25 ml LB, incubated overnight at 37°C, 250 rpm, and subsequently inoculated into 200 ml LB supplemented with 50 µM $ZnCl_2$ for an OD at 600 nm of 0.02. Incubated at 37°C, 250 rpm and induced expression by adding 100 µM IPTG. Incubated for further 6h. Centrifuged at 4K rpm and lysed with BC-500 (20 mM Tri-HCl pH 7.4, 2 mM EDTA, 20% glycerol, 500 mM KCl, 1 mM DTT and 0.5 mM PMSF). Sonicated on ice (10X 10 sec ON, 10 sec OFF). Added NP-40 to a final concentration of 0.1% and incubated for 30 minutes at 4°C, top over end. Centrifuged at 14K rpm for 10 minutes at 4°C, saved the supernatant and stored at -80°C.
For GST fusion protein purification, thawed one aliquot of protein lysate and added 50 µl Glutathione Sepharose 4B beads (50% slurry, equilibrated with BC-500) and incubated overnight at 4°C, top over end. Washed the beads 5 times with BC-500 with 0.1% NP-40 and two additional washes (20 minutes at RT, top over end) with BC-180 (same as BC-500 but with 180 mM KCl) supplemented with 0.2% sarcosyl. Ressuspended the beads in 40 µl laemmli buffer, denatured for 10 minutes at 80°C and resolved in a bis-tris 4-12% gel, together with BSA standards to determine purified protein concentration. These concentrations were used to adjust the starting amount of lysate for subsequent IPs.
For each IP, the adequate amount of lysate to yield 1 µg of GST-fusion was thawed and processed as described above. After the final wash, beads were incubated with 1 µg total RNA (isolated from mouse liver) in BC-180 supplemented with 0.05% NP-40 for 1h at RT, half rotation top over end. Beads were washed 5 times with BC-180 supplemented with 0.0% NP-40, followed by 3 elutions in 100 µl of 1% SDS, 0.1 M $NaHCO_3$. Added 12 µl of 5M NaCl and 6 µl Proteinase K (Qiagen). Incubated overnight at 62°C followed by 5 minutes at 95°C. RNA was ethanol precipitated in the presence of 50 µg glycogen carrier. Incubated 30 minutes at RT followed by 1h at -20°C. Centrifuged for 20 minutes at 14K rpm at 4°C, washed twice with 70% ethanol, dried and finally ressuspended in 20 µl nuclease-free water and quantified using a nanodrop.

**Publicly available datasets**

Data for *Zfp697/ZNF697* expression in different animal models and human cohorts were collected from the following studies deposited on GEO database: GSE211204 (human skeletal muscle; unilateral limb suspension) (*21*); GSE210263 (mouse skeletal muscle; muscular dystrophy with myositis, mdm) (*22*); GSE114820 (mouse skeletal muscle; Cancer Cachexia) (*23*); GSE202295 (human skeletal muscle; type 2 diabetes) (*24*); GSE98622 (mouse kidney; acute kidney injury) (*25*); GSE153494 (mouse heart; myocardial infarction) (*26*); GSE198926 (mouse heart; transverse aortic constriction) (*27*); GSE151834 (mouse heart; Ischemic cardiac injury) (*28*); GSE147127 (nuclei populations in mouse soleus and tibialis anterior, single-nuclei RNA sequencing (*29*); GSE143437 (mouse mononuclear cells, single-cell RNA sequencing) (*15*). Data for *ZNF697* response following acute exercise in humans were retrieved from the MetaMex database (*30*).

**Quantification and statistical analysis**

The analysis of the data was carried out using R, Excel, and Prism as described above. Specifics regarding the statistical analyses and sample sizes are found in the Figure Legends. Outliers were detected using Grubb's test and removed when $P<0.05$.

### Supplementary Figures

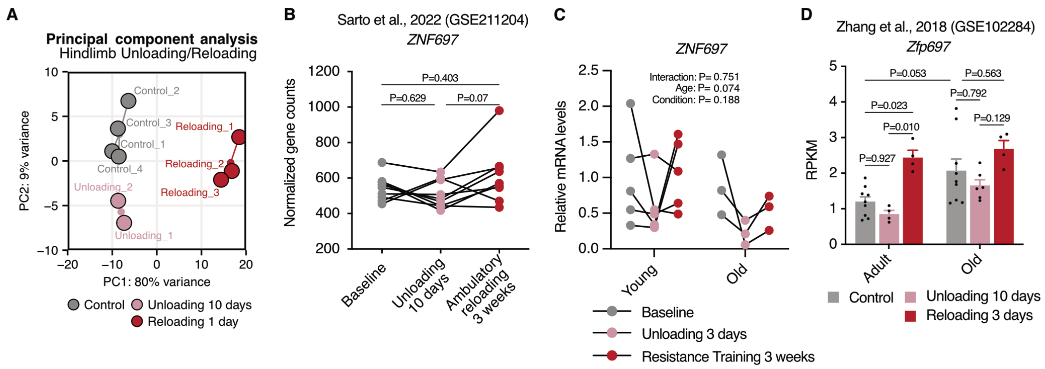


**Fig. S1. Zfp697/ZNF697 expression is increased in mouse and human muscle during reloading.** (**A**) Principal component analysis for the RNA-seq performed in the gastrocnemius muscle of mice after 10 days of hindlimb unloading, 10 days of unloading followed by 1 day of reloading, and controls (n = 3 per condition). (**B**) *ZNF697* gene expression in human muscle biopsies collected subjects at baseline, after 10 days of limb suspension and after 3 weeks of ambulatory recovery. Publicly available dataset (GSE211204) (*21*). **(C)** *ZNF697* gene expression in human muscle biopsies collected subjects at baseline, after 3 days of limb suspension and after 3 weeks of resistance training. (**D**) *Zfp697* gene expression in soleus muscle of adult (6-month-old) and old (22- to 24-month-old) mice after 10 days of hindlimb unloading, 3 days of reloading, and controls (n = 4-9 per condition). Publicly available dataset (GSE102284) (*34*)(*31*). Two-way ANOVA with Tukey's multiple comparisons test. Data represent mean values and error bars represent SEM.

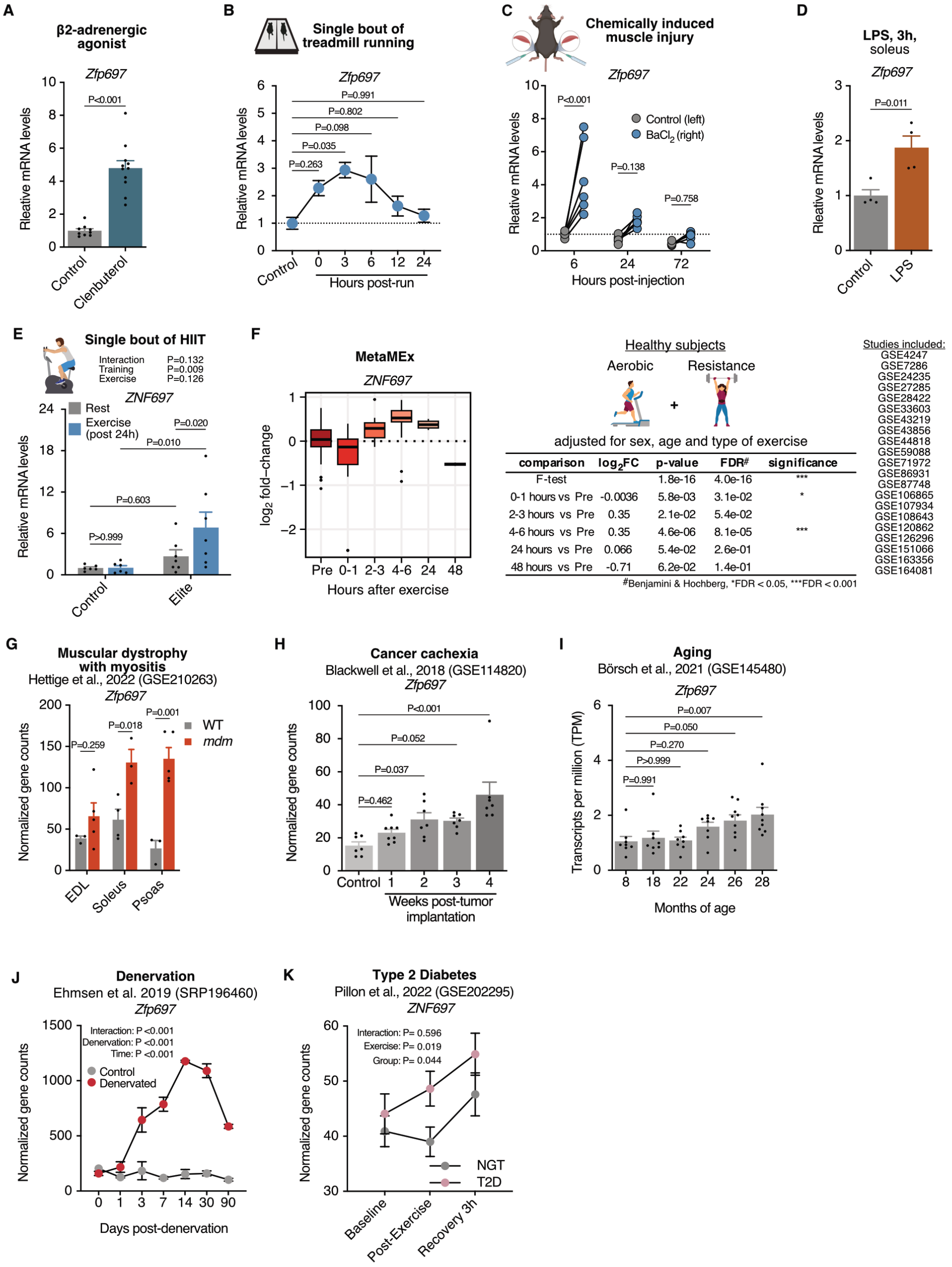


**Fig. S2. Zfp697/ZNF697 expression is transiently elevated during recovery from injury.** (**A**) *Zfp697* gene expression in mouse gastrocnemius after 16 hours of clenbuterol injection (i.p. 2 mg/kg, n = 9-11). Two-tailed Student's t test. (**B**) *Zfp697* gene expression in mouse quadriceps following a single bout of treadmill running at 0, 3, 6, 12 and 24 hours, compared with the rest control (n = 4-6). One-way ANOVA with Dunnett's multiple comparisons test. (**C**) *Zfp697* gene expression in mouse gastrocnemius after chemically induced muscle injury. $BaCl_2$ was intramuscularly injected in the gastrocnemius of one leg (right) and control solution in the same site of contralateral leg (left). Muscles were harvested 6, 24 and 72 hours post-injection (n = 6 per time point). Two-way ANOVA with Šídák's multiple comparisons test. (**D**) *Zfp697* gene expression in mouse soleus after 3 hours of LPS injection (i.p. 1 g/g, n = 4). Two-tailed Student's t test. (**E**) *ZNF697* gene expression in muscle biopsies taken before and 24 hours after a single bout of high-intensity interval training in recreationally active subjects (Control) and elite athletes (Elite) (n = 6-7). Two-way ANOVA with Šídák's multiple comparisons test. (**F**) *ZNF697* gene expression in human skeletal muscle after exercise. Data collected from the MetaMEx database (*30*). (**G**) *Zfp697* gene expression in skeletal muscles of mice with muscular dystrophy with myositis (mdm) compared with wild-type (WT) controls. Publicly available dataset (GSE210263) (*22*). Two-tailed Student's t test. (**H**) *Zfp697* gene expression in mouse gastrocnemius muscle throughout the progression of cancer cachexia. Publicly available dataset (GSE114820) (*23*). One-way ANOVA with Dunnett's multiple comparisons test. (**I**) *Zfp697* gene expression in mouse gastrocnemius muscle during aging. Publicly available dataset (GSE145480) (*32*). One-way ANOVA with Dunnett's multiple comparisons test. (**J**) *Zfp697* gene expression in mouse gastrocnemius muscle in response to denervation injury compared to contralateral control muscle. Publicly available dataset (SRP196460) (*33*). Repeated measures two-way ANOVA. (**K**) *ZNF697* gene expression in muscle biopsies collected after endurance exercise in normoglycemic (NGT) and type 2 diabetic patients (T2D). Publicly available dataset (GSE202295) (*24*). Data represent mean values and error bars represent SEM.

### Figure S3

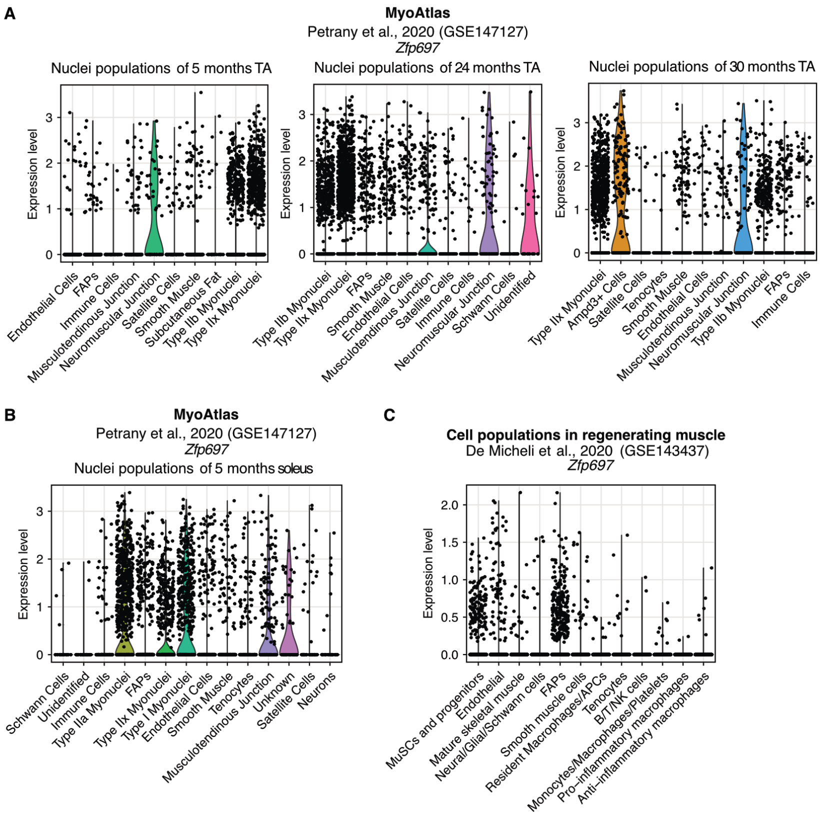


**Fig. S3. Zfp697 is expressed both in mature muscle fibers and muscle-resident cell populations.** (**A**) *Zfp697* gene expression in nuclei isolated from tibialis anterior (TA) of 5-, 24- and 30-month-old mice. Publicly available dataset (GSE147127) (*29*). (**B**) *Zfp697* gene expression in nuclei isolated from soleus of 5-month-old mice. Publicly available dataset (GSE147127) (*29*). (**C**) *Zfp697* gene expression in mononuclear cells isolated from regenerating skeletal muscle. Publicly available dataset (GSE143437) (*15*).

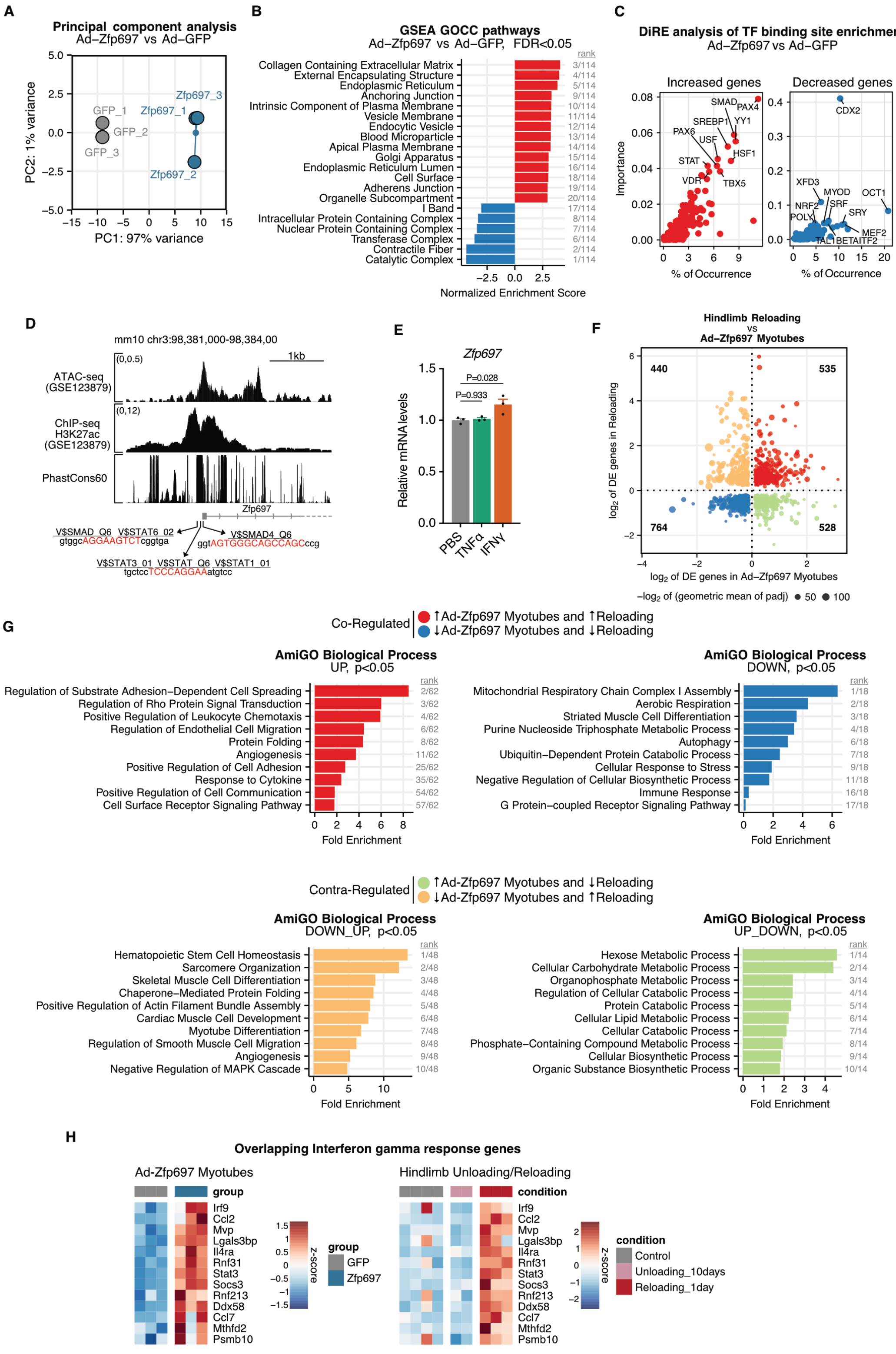


**Fig. S4. Genes in common between Zfp697 expression in myotubes and muscle reloading highlight IFNγ response.** (**A**) Principal component analysis for RNA-seq performed in mouse primary myotubes overexpressing Zfp697. (**B**) Gene set enrichment analysis for gene ontology cellular components (GOCC) in mouse primary myotubes overexpressing Zfp697. FDR, false discovery rate. (**C**) DiRE analysis for enriched transcription factor binding sites in the upstream regulatory regions of genes increased or decreased by Zfp697 overexpression in mouse primary myotubes. (**D**) Genome browser tracks of the mouse *Zfp697* promoter highlighting skeletal muscle ATAC-seq and H3K27ac ChIP-seq (GSE123879) (*34*), and PhastCons60 vertebrate conservation score. Predicted transcription factor binding sites were identified using rVista2.0 (*35*). (**E**) *Zfp697* gene expression in mouse primary myotubes treated with vehicle (PBS), recombinant tumor necrosis factor alpha (TNFα; 200 ng/mL) or interferon gamma (IFNγ; 20 ng/mL) for 8 hours. One-way ANOVA with Dunnett's multiple comparisons test. Data represent mean values and error bars represent SEM. (**F**) Correlation plot for differentially expressed (DE, fold-change) genes in gastrocnemius muscle following 1 day of hindlimb reloading compared with 10 days of hindlimb unloading, and mouse primary myotubes overexpressing Zfp697. padj, adjusted p-value. (**G**) Top gene ontology biological processes overrepresented in the genes co-regulated and contra-regulated from Figure S3F. (**H**) Heatmap for interferon gamma response genes commonly increased during hindlimb reloading in muscle and Zfp697 overexpression in primary myotubes.

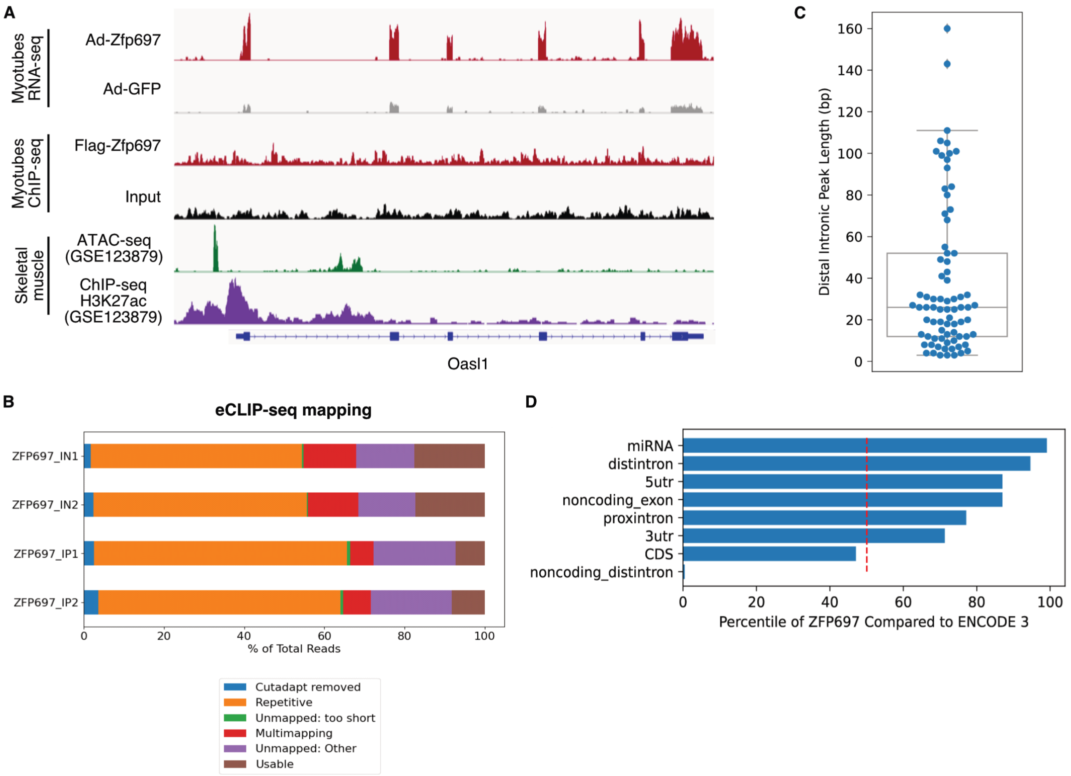


**Fig. S5. Zfp697 is not primarily a DNA-binding transcription factor.** (**A**) Genome browser tracks of the mouse *Oasl1* gene showing RNA-seq data from mouse primary myotubes overexpressing Zfp697 and ChIP-seq data for Flag-tagged Zfp697, also highlighting skeletal muscle ATAC-seq and H3K27ac ChIP-seq (GSE123879) (*34*). (**B**) Mapping of Zfp697 eCLIP and input reads. (**C**) Length of Zfp697 enriched peaks (binding sites) mapped to distal introns. (**D**) Percentiles of the proportion of Zfp697 binding sites mapped to various genic regions compared to RBPs with eCLIP data in ENCODE 3.

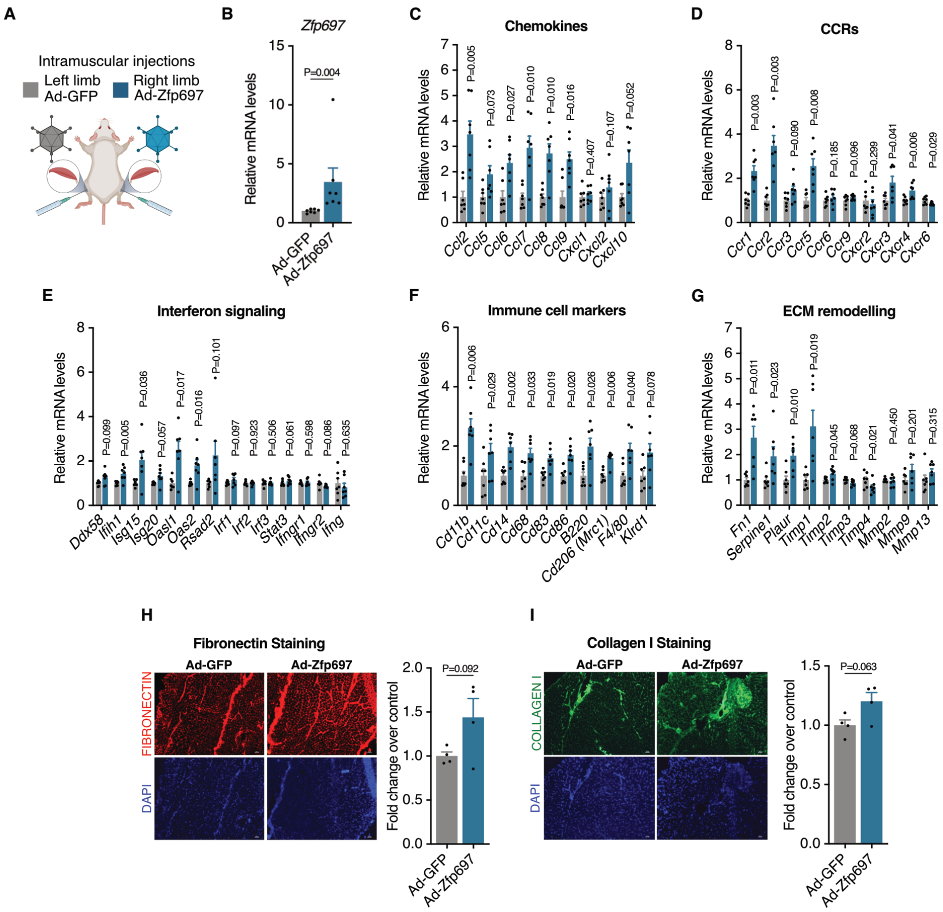


**Fig. S6. Sustained Zfp697 expression *in vivo* increases expression of inflammation and ECM remodeling.** (**A**) Schematic representation of adenovirus-mediated expression of Zfp697 in mouse skeletal muscle. Each gastrocnemius of 14-day-old SCID mice was injected with adenovirus expressing GFP alone or with Zfp697 and harvested after 7 days. (**B-G**) Gene expression analysis of *Zfp697*, chemokines, chemokine receptors (CCRs), interferon signaling, immune cell markers and extracellular matrix (ECM) remodelling markers in the gastrocnemius muscle of mice intramuscularly injected with Ad-Zfp697 or Ad-GFP (n = 6). Two-tailed Student's paired t test versus Ad-GFP. (**H-I**) Representative immunostainings for fibronectin (H) and collagen I (I), and respective quantification, in the gastrocnemius muscle of mice intramuscularly injected with Ad-Zfp697 or Ad-GFP (n = 4). DAPI was used to stain nuclei. Two-tailed Student's paired t test. Data represent mean values and error bars represent SEM.

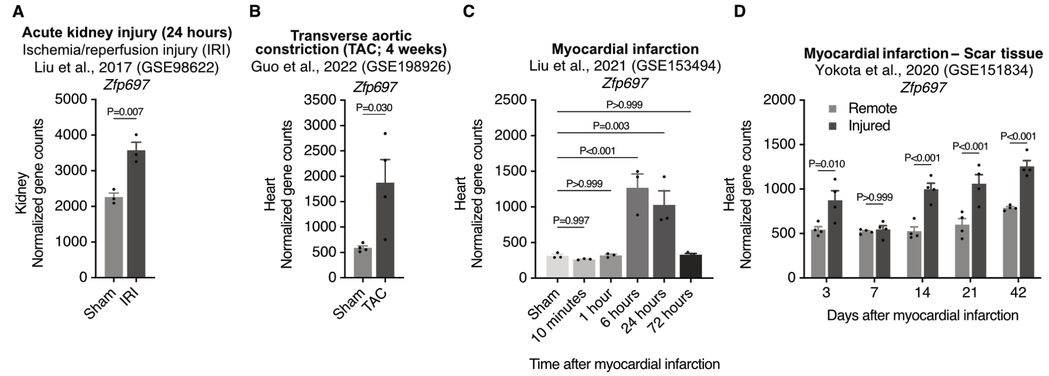


**Fig. S7. Zfp697 expression in different metabolic tissues upon injury/disease**. (**A**) *Zfp697* gene expression in mouse kidney 24 hours after acute kidney injury by ischemia reperfusion injury (IRI) or sham surgery. Publicly available dataset (GSE98622) (*25*). Two-tailed Student's t test. (**B**) *Zfp697* gene expression in mouse heart 4 weeks after transverse aortic constriction (TAC) or sham surgery. Publicly available dataset (GSE198926) (*27*). Two-tailed Student's t test. (**C**) *Zfp697* gene expression in mouse heart in response to myocardial infarction compared to sham surgery. Publicly available dataset (GSE153494) (*26*). One-way ANOVA with Dunnett's multiple comparisons test. (**D**) *Zfp697* gene expression in injured myocardium (scar tissue) compared to remote myocardium region in response to myocardial infarction in mice. Publicly available dataset (GSE151834) (*28*). Two-way ANOVA with Šídák's multiple comparisons test. Data represent mean values and error bars represent SEM.

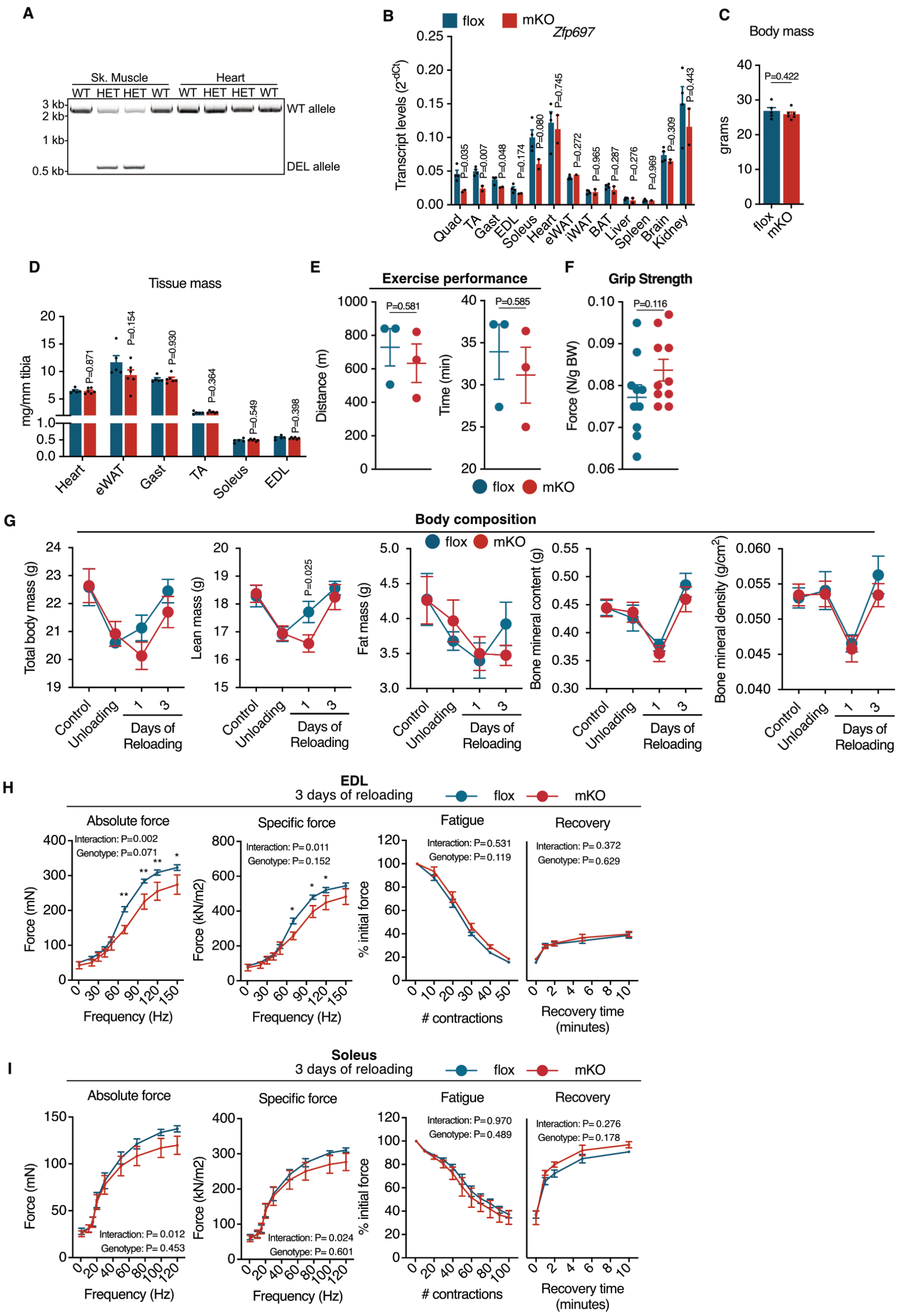


**Fig. S8. Loss of Zfp697 in muscle fibers leads to reduced muscle force.** (**A**) PCR amplification of the deleted segment of exon 3 in skeletal and heart muscle of Zfp697-mKO heterozygous mice. (**B**) Tissue panel for Zfp697 gene expression in Zfp697-mKO and flox littermates (n = 2-3). Two-tailed Student's t test. (**C**) Body mass of Zfp697-mKO and flox littermates (n = 5-6). Two-tailed Student's t test. (**D**) Tissue mass of heart, epididymal white adipose tissue (eWAT), gastrocnemius (Gast), tibialis anterior (TA), soleus and extensor digitorium longus (EDL) Zfp697-mKO and flox littermates, normalized by tibia length (n = 5-6). Two-tailed Student's t test. (**E-F**) Total distance and time achieved in a treadmill running to exhaustion test (left panel; n = 3) and body-weight-normalized grip strength test (right panel; n = 10) of Zfp697-mKO and flox littermates. Two-tailed

Student's t test. (**G**) Analysis of body composition by dual-energy X-ray absorptiometry in Zfp697-mKO and flox littermates subjected to hindlimb unloading and reloading (n = 4-8). Two-way ANOVA with Fisher's LSD. P values represent comparison with time-matched flox controls. (**H-I**) *Ex vivo* analysis of contractile force, fatigability, and recovery of EDL (H) and soleus (I) muscles from Zfp697-mKO and floxed littermates subjected to hindlimb unloading followed by 3 days of reloading (n = 4-6). Repeated measures two-way ANOVA with Fisher's LSD. P values represent comparison with frequency-matched flox controls. *P<0.05, **P<0.001. Data represent mean values and error bars represent SEM.

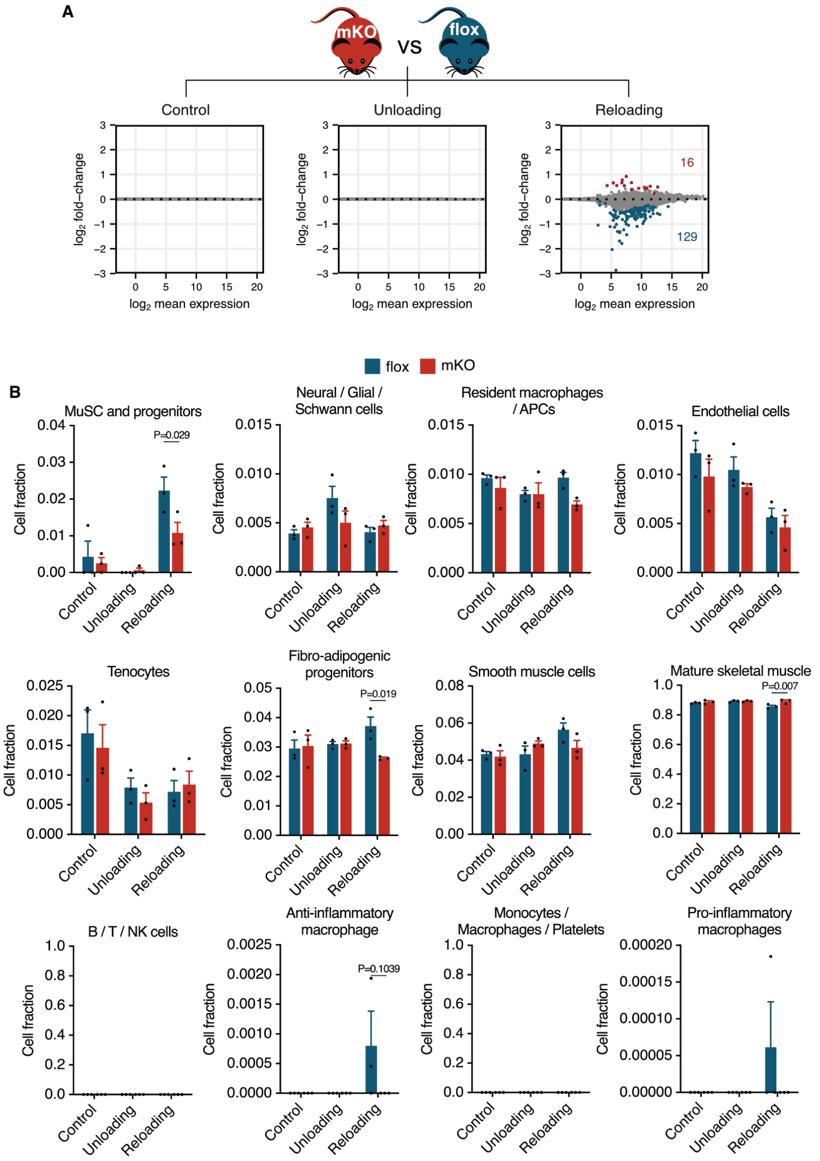


**Fig. S9. Loss of muscle fiber Zfp697 prevents satellite cell and FAP activation.** (**A**) MA Plots for RNA-seq performed in the gastrocnemius muscle of control Zfp697 flox and mKO, after 10 days of hindlimb unloading and 3 days of reloading (n = 3 per condition and genotype). Red dots indicate genes with significantly increased expression, while blue dots indicate genes with significantly decreased expression (FDR < 0.05), accounting for the interaction effect between conditions (control, unloading and reloading) and genotypes (mKO and flox). FDR, false discovery rate. (**B**) Muscle-resident cell populations identified by digital cytometry (CIBERSORTx) applied to bulk muscle RNA-seq data from control, hindlimb unloaded and reloaded Zfp697 flox and mKO mice (n=3 per condition and genotype). Two-way ANOVA with Šídák’s multiple comparisons test. Data represent mean values and error bars represent SEM.

### References and Notes


1. E. R. Morey-Holton, R. K. Globus, Hindlimb unloading rodent model: technical aspects. *Journal of applied physiology (Bethesda, Md. : 1985)*. **92**, 1367–77 (2002).

2. T. a Rando, H. M. Blau, Primary mouse myoblast purification, characterization, and transplantation for cell-mediated gene therapy. *The Journal of cell biology*. **125**, 1275–87 (1994).

3. S. Li, M. P. Czubryt, J. McAnally, R. Bassel-Duby, J. A. Richardson, F. F. Wiebel, A. Nordheim, E. N. Olson, Requirement for serum response factor for skeletal muscle growth and maturation revealed by tissue-specific gene deletion in mice. *Proc. Natl. Acad. Sci. U.S.A.* **102**, 1082–1087 (2005).

4. N. Place, N. Ivarsson, T. Venckunas, D. Neyroud, M. Brazaitis, A. J. Cheng, J. Ochala, S. Kamandulis, S. Girard, G. Volungevičius, H. Paužas, A. Mekideche, B. Kayser, V. Martinez-Redondo, J. L. Ruas, J. Bruton, A. Truffert, J. T. Lanner, A. Skurvydas, H. Westerblad, Ryanodine receptor fragmentation and sarcoplasmic reticulum $Ca^{2+}$ leak after one session of high-intensity interval exercise. *Proc. Natl. Acad. Sci. U.S.A.* **112**, 15492–15497 (2015).

5. T. Venckunas, M. Brazaitis, A. Snieckus, M. Mickevicius, N. Eimantas, A. Subocius, D. Mickeviciene, H. Westerblad, S. Kamandulis, Adding High-Intensity Interval Training to Classical Resistance Training Does Not Impede the Recovery from Inactivity-Induced Leg Muscle Weakness. *Antioxidants*. **12**, 16 (2023).

6. S. K. Akula, K. B. McCullough, C. Weichselbaum, J. D. Dougherty, S. E. Maloney, The trajectory of gait development in mice. *Brain Behav*. **10** (2020), doi:10.1002/brb3.1636.

7. A. Dobin, C. A. Davis, F. Schlesinger, J. Drenkow, C. Zaleski, S. Jha, P. Batut, M. Chaisson, T. R. Gingeras, STAR: ultrafast universal RNA-seq aligner. *Bioinformatics*. **29**, 15–21 (2013).

8. Y. Liao, G. K. Smyth, W. Shi, featureCounts: an efficient general purpose program for assigning sequence reads to genomic features. *Bioinformatics*. **30**, 923–930 (2014).

9. M. I. Love, W. Huber, S. Anders, Moderated estimation of fold change and dispersion for RNA-seq data with DESeq2. *Genome Biol*. **15**, 550 (2014).

10. M. Stephens, False discovery rates: a new deal. *Biostat*, kxw041 (2016).

11. A. Subramanian, P. Tamayo, V. K. Mootha, S. Mukherjee, B. L. Ebert, M. A. Gillette, A. Paulovich, S. L. Pomeroy, T. R. Golub, E. S. Lander, J. P. Mesirov, Gene set enrichment analysis: A knowledge-based approach for interpreting genome-wide expression profiles. *Proceedings of the National Academy of Sciences*. **102**, 15545–15550 (2005).

12. A. Liberzon, C. Birger, H. Thorvaldsdóttir, M. Ghandi, J. P. Mesirov, P. Tamayo, The Molecular Signatures Database Hallmark Gene Set Collection. *cels*. **1**, 417–425 (2015).

13. S. Carbon, A. Ireland, C. J. Mungall, S. Shu, B. Marshall, S. Lewis, AmiGO: online access to ontology and annotation data. *Bioinformatics*. **25**, 288–289 (2009).

14. A. M. Newman, C. B. Steen, C. L. Liu, A. J. Gentles, A. A. Chaudhuri, F. Scherer, M. S. Khodadoust, M. S. Esfahani, B. A. Luca, D. Steiner, M. Diehn, A. A. Alizadeh, Determining cell type abundance and expression from bulk tissues with digital cytometry. *Nat Biotechnol*. **37**, 773–782 (2019).

15. A. J. De Micheli, E. J. Laurilliard, C. L. Heinke, H. Ravichandran, P. Fraczek, S. Soueid-Baumgarten, I. De Vlaminck, O. Elemento, B. D. Cosgrove, Single-Cell Analysis of the Muscle Stem Cell Hierarchy Identifies Heterotypic Communication Signals Involved in Skeletal Muscle Regeneration. *Cell Rep*. **30**, 3583-3595.e5 (2020).

16. J. Feng, T. Liu, B. Qin, Y. Zhang, X. S. Liu, Identifying ChIP-seq enrichment using MACS. *Nat Protoc*. **7**, 1728–1740 (2012).

17. A. R. Quinlan, I. M. Hall, BEDTools: a flexible suite of utilities for comparing genomic features. *Bioinformatics*. **26**, 841–842 (2010).

18. E. L. Van Nostrand, G. A. Pratt, A. A. Shishkin, C. Gelboin-Burkhart, M. Y. Fang, B. Sundararaman, S. M. Blue, T. B. Nguyen, C. Surka, K. Elkins, R. Stanton, F. Rigo, M. Guttman, G. W. Yeo, Robust transcriptome-wide discovery of RNA-binding protein binding sites with enhanced CLIP (eCLIP). *Nat Methods*. **13**, 508–514 (2016).

19. M. T. Lovci, D. Ghanem, H. Marr, J. Arnold, S. Gee, M. Parra, T. Y. Liang, T. J. Stark, L. T. Gehman, S. Hoon, K. B. Massirer, G. A. Pratt, D. L. Black, J. W. Gray, J. G. Conboy, G. W. Yeo, Rbfox proteins regulate alternative mRNA splicing through evolutionarily conserved RNA bridges. *Nat Struct Mol Biol*. **20**, 1434–1442 (2013).

20. J. Luo, Z.-L. Deng, X. Luo, N. Tang, W.-X. Song, J. Chen, K. A. Sharff, H. H. Luu, R. C. Haydon, K. W. Kinzler, B. Vogelstein, T.-C. He, A protocol for rapid generation of recombinant adenoviruses using the AdEasy system. *Nat Protoc*. **2**, 1236–1247 (2007).

21. F. Sarto, D. W. Stashuk, M. V. Franchi, E. Monti, S. Zampieri, G. Valli, G. Sirago, J. Candia, L. M. Hartnell, M. Paganini, J. S. McPhee, G. De Vito, L. Ferrucci, C. Reggiani, M. V. Narici, Effects of short-term unloading and active recovery on human motor unit properties, neuromuscular junction transmission and transcriptomic profile. *The Journal of Physiology*. **n/a** (2022), doi:10.1113/JP283381.

22. P. Hettige, U. Tahir, K. C. Nishikawa, M. J. Gage, Transcriptomic profiles of muscular dystrophy with myositis (mdm) in extensor digitorum longus, psoas, and soleus muscles from mice. *BMC Genomics*. **23**, 657 (2022).

23. T. A. Blackwell, I. Cervenka, B. Khatri, J. L. Brown, M. E. Rosa-Caldwell, D. E. Lee, R. A. Perry, L. A. Brown, W. S. Haynie, M. P. Wiggs, W. G. Bottje, T. A. Washington, B. C. Kong, J. L. Ruas, N. P. Greene, Transcriptomic analysis of the development of skeletal muscle atrophy in cancer-cachexia in tumor-bearing mice. *Physiological Genomics*. **50**, 1071–1082 (2018).

24. N. J. Pillon, J. A. B. Smith, P. S. Alm, A. V. Chibalin, J. Alhusen, E. Arner, P. Carninci, T. Fritz, J. Otten, T. Olsson, S. van Doorslaer de ten Ryen, L. Deldicque, K. Caidahl, H. Wallberg-Henriksson, A. Krook, J. R. Zierath, Distinctive exercise-induced inflammatory response and exerkine induction in skeletal muscle of people with type 2 diabetes. *Science Advances*. **8**, eabo3192 (2022).

25. J. Liu, S. Kumar, E. Dolzhenko, G. F. Alvarado, J. Guo, C. Lu, Y. Chen, M. Li, M. C. Dessing, R. K. Parvez, P. E. Cippà, A. M. Krautzberger, G. Saribekyan, A. D. Smith, A. P. McMahon, Molecular characterization of the transition from acute to chronic kidney injury following ischemia/reperfusion. *JCI Insight*. **2**, e94716, 94716 (2017).

26. W. Liu, J. Shen, Y. Li, J. Wu, X. Luo, Y. Yu, Y. Zhang, L. Gu, X. Zhang, C. Jiang, J. Li, Pyroptosis inhibition improves the symptom of acute myocardial infarction. *Cell Death Dis*. **12**, 852 (2021).

27. A. H. Guo, R. Baliira, M. E. Skinner, S. Kumar, A. Andren, L. Zhang, R. S. Goldsmith, S. Michan, N. J. Davis, M. W. Maccani, S. M. Day, D. A. Sinclair, M. J. Brody, C. A. Lyssiotis, A. B. Stein, D. B. Lombard, Sirtuin 5 levels are limiting in preserving cardiac function and suppressing fibrosis in response to pressure overload. *Sci Rep*. **12**, 12258 (2022).

28. T. Yokota, J. McCourt, F. Ma, S. Ren, S. Li, T.-H. Kim, Y. Z. Kurmangaliyev, R. Nasiri, S. Ahadian, T. Nguyen, X. H. M. Tan, Y. Zhou, R. Wu, A. Rodriguez, W. Cohn, Y. Wang, J. Whitelegge, S. Ryazantsev, A. Khademhosseini, M. A. Teitell, P.-Y. Chiou, D. E. Birk, A. C. Rowat, R. H. Crosbie, M. Pellegrini, M. Seldin, A. J. Lusis, A. Deb, Type V Collagen in Scar Tissue Regulates the Size of Scar after Heart Injury. *Cell*. **182**, 545-562.e23 (2020).

29. M. J. Petrany, C. O. Swoboda, C. Sun, K. Chetal, X. Chen, M. T. Weirauch, N. Salomonis, D. P. Millay, Single-nucleus RNA-seq identifies transcriptional heterogeneity in multinucleated skeletal myofibers. *Nat Commun*. **11**, 6374 (2020).

30. N. J. Pillon, B. M. Gabriel, L. Dollet, J. A. B. Smith, L. Sardón Puig, J. Botella, D. J. Bishop, A. Krook, J. R. Zierath, Transcriptomic profiling of skeletal muscle adaptations to exercise and inactivity. *Nat Commun*. **11**, 470 (2020).

31. X. Zhang, M. B. Trevino, M. Wang, S. J. Gardell, J. E. Ayala, X. Han, D. P. Kelly, B. H. Goodpaster, R. B. Vega, P. M. Coen, Impaired mitochondrial energetics characterize poor early recovery of muscle mass following hind limb unloading in old mice. *Journals of Gerontology - Series A Biological Sciences and Medical Sciences*. **73**, 1313–1322 (2018).

32. A. Börsch, D. J. Ham, N. Mittal, L. A. Tintignac, E. Migliavacca, J. N. Feige, M. A. Rüegg, M. Zavolan, Molecular and phenotypic analysis of rodent models reveals conserved and species-specific modulators of human sarcopenia. *Commun Biol*. **4**, 1–15 (2021).

33. J. T. Ehmsen, R. Kawaguchi, R. Mi, G. Coppola, A. Höke, Longitudinal RNA-Seq analysis of acute and chronic neurogenic skeletal muscle atrophy. *Sci Data*. **6**, 179 (2019).

34. K. Ramachandran, M. D. Senagolage, M. A. Sommars, C. R. Futtner, Y. Omura, A. L. Allred, G. D. Barish, Dynamic enhancers control skeletal muscle identity and reprogramming. *PLoS biology*. **17**, e3000467 (2019).

35. G. G. Loots, I. Ovcharenko, rVISTA 2.0: evolutionary analysis of transcription factor binding sites. *Nucleic Acids Res*. **32**, W217–W221 (2004).

**Data S1.** RNA-Seq analysis of mouse gastrocnemius muscle after 10 days of hindlimb unloading, 10 days of unloading followed by 1 day of reloading, and controls. **(separate file)**.

**Data S2.** GSEA for hallmark pathways in mouse gastrocnemius after hindlimb unloading and reloading. **(separate file)**

**Data S3.** RNA-Seq analysis of primary mouse myotubes overexpressing Zfp967. **(separate file)**

**Data S4.** GSEA for hallmark pathways and GOCC of primary mouse myotubes overexpressing Zfp967. **(separate file)**

**Data S5.** Genes co-regulated/contra-regulated and gene ontology analysis for hindlimb reloading compared to mouse primary myotubes overexpressing Zfp697. **(separate file)**

**Data S6.** Zfp697 eCLIP peaks in mouse primary myotubes overexpressing Zfp697. **(separate file)**

**Data S7.** RNA-seq analysis of gastrocnemius muscle from control Zfp697 flox and mKO mice, after 10 days of hindlimb unloading and 3 days of reloading. **(separate file)**

**Data S8.** GSEA for hallmark pathways of gastrocnemius muscle from Zfp697 flox and mKO in response to hindlimb reloading compared to unloading. **(separate file)**
